# Accounting for unobserved population dynamics and aging error in close-kin mark-recapture assessments

**DOI:** 10.1101/2023.02.20.529265

**Authors:** John D. Swenson, Elizabeth N. Brooks, Dovi Kacev, Charlotte Boyd, Michael Kinney, Benjamin Marcy-Quay, Anthony Sévêque, Kevin Feldheim, Lisa M. Komoroske

## Abstract

Obtaining robust estimates of population abundance is a central challenge hindering the conservation and management of many threatened and exploited species. Close-kin mark-recapture (CKMR) is a genetics-based approach that has strong potential to improve monitoring of data-limited species by enabling estimates of abundance, survival, and other parameters for populations that are challenging to assess. However, CKMR models have received limited sensitivity testing under realistic population dynamics and sampling scenarios, impeding application of the method in population monitoring programs and stock assessments. Here, we use individual-based simulation to examine how unmodeled population dynamics and aging uncertainty affect the accuracy and precision of CKMR parameter estimates under different sampling strategies. We then present adapted models that correct the biases that arise from model misspecification. Our results demonstrate that a simple base-case CKMR model produces robust estimates of population abundance with stable populations that breed annually; however, if a population trend or non-annual breeding dynamics are present, or if year-specific estimates of abundance are desired, a more complex CKMR model must be constructed. In addition, we show that CKMR can generate reliable abundance estimates for adults from a variety of sampling strategies, including juvenile-focused sampling where adults are never directly observed (and aging error is minimal). Finally, we apply a CKMR model that has been adapted for population growth and intermittent breeding to two decades of genetic data from juvenile lemon sharks (*Negaprion brevirostris*) in Bimini, Bahamas, to demonstrate how application of CKMR to samples drawn solely from juveniles can contribute to monitoring efforts for highly mobile populations. Overall, this study expands our understanding of the biological factors and sampling decisions that cause bias in CKMR models, identifies key areas for future inquiry, and provides recommendations that can aid biologists in planning and implementing an effective CKMR study, particularly for long-lived data-limited species.

## Introduction

Population abundance plays important roles in both fundamental and applied biological research and is associated with a wide range of ecological and evolutionary processes (Hassell 1975, Berryman 1989, Robertson 1996, Carbone et al. 2011, Ellegren and Galtier 2016). Abundance estimates and trends are also key metrics for conservation and management and are commonly used to assess conservation status (Wilson et al. 2011), quantify the impacts of threats and/or recovery efforts (Jennings 2000, Ward-Paige et al. 2012, Magera et al. 2013), and scale regulated harvest quantities (e.g., allowable biological catch, annual catch limits) for managed populations of target and non-target species. Consequently, a wide range of methods have been developed for estimating population abundance (Schwarz and Seber 1999, Wilson and Delahay 2001, McCauley et al. 2012).

Capture-mark-recapture (CMR) is one prominent and widely used method in which abundance is estimated by constructing capture histories for each sampled (or tagged) individual, estimating capture probabilities, and comparing the number of recaptured individuals to the total number of sampled individuals (Cormack 1964, Jolly 1965, Seber 1965). A number of variations of CMR methods have been developed over the years to account for varied population demographics and sampling schemes (Pollock 2000, Amstrup et al. 2010, Royle et al. 2013), but the approach remains largely intractable in situations where recapture rates are very low, as with many low density and highly mobile marine species (Kohler and Turner 2001, Webster et al. 2002, Boyd et al. 2018). In addition, estimating a capture probability for CMR requires an estimate of the rate at which tags are lost (Arnason and Mills 1981, Hyun et al. 2012) and reported (e.g., by fishermen or hunters; Green et al. 1983, Pollock et al. 2001, Sackett and Catalano 2017), and tag loss and reporting rates vary with the species and experimental design (Oosthuizen et al. 2010). As such, their estimation is likely to require auxiliary studies that demand more time and resources and may be reliant on cooperation from individuals that encounter the tags. CMR provides direct information about the sampled demographic, but many highly mobile marine species have spatially segregated life histories and are only available for sampling in nearshore habitats as juveniles before transitioning to a less accessible pelagic habitat as adults. In such cases, CMR results are restricted to providing direct information about the juvenile portion of the population, while the population dynamics of adults can only be modeled effectively if additional data are available and if key assumptions are met (Kendall 1999, Pollock 2000). As alternatives to CMR, surveys or transect-based methods can be helpful tools to estimate regional abundance (Schwarz and Seber 1999). However, variability in survey length, uncertainty surrounding the proportion of habitat sampled and shifts in habitat availability, as well as changes in behavior arising from the presence of human observers and observation error are common pitfalls that can make such methods unreliable or incomparable across studies (McCauley et al. 2012, Boyd and Punt 2021, Davis et al. 2022).

While CMR, surveys, and transect-based methods can all be useful tools for abundance estimation in certain contexts, applying them in an unbiased way can be prohibitively challenging in many systems. When estimates of absolute abundance are infeasible, indices of relative abundance are commonly used to assess populations of exploited species (Campbell 2015). In fisheries, abundance trends derived from catch and effort data (e.g., catch-per-unit-effort, CPUE), in concert with biological reference points, can inform management by providing critical information about whether a population is overfished or if overfishing is actively occurring (Cortés and Brooks 2018). However, it is extremely challenging to account for all the factors that could influence catchability (Maunder et al. 2006); hence, indices of relative abundance derived from CPUE are rarely linearly proportional to actual abundance (Harley et al. 2001, Maunder and Punt 2004, Lynch et al. 2012). Fish or fisher behavior contributes to hyperstability (Erisman et al. 2011, Ward et al. 2013) and biased inference about abundance trends can result if CPUE data are interpreted in isolation, or if linearity between catch rate and abundance is implicitly assumed (Maunder et al. 2006). Furthermore, estimating trends of relative abundance for highly mobile species frequently requires the integration of multiple independent surveys that suggest differing abundance trends, making it difficult to establish true abundance patterns (Peterson et al. 2021). All of these issues are amplified in taxa such as elasmobranchs (sharks, skates, and rays), where reported catch data are often unreliable (Cortés and Brooks 2018). While CPUE can provide invaluable information regarding stock status and harvest pressure when analyzed in the right context (e.g., via an integrated model that incorporates additional data streams), there is an urgent need for methods that can provide robust estimates of absolute population abundance in circumstances where catch data are unreliable or strongly correlated with factors other than population trend (e.g., changes in fishing practices, skill, or gear improvement, or environmental perturbations).

Close-kin mark-recapture (CKMR) is a genetics-based approach for estimating absolute population abundance that overcomes many of the logistical challenges associated with CMR and other abundance estimation methods (Skaug 2001, Bravington et al. 2016a). As such, CKMR has great potential to expand monitoring efforts and improve or enable assessments of species for which conventional methods are intractable. In contrast to conventional CMR, the tags in CKMR are genotypes, and animals are considered “re-captured” when their kin are identified (Bravington et al. 2016a). This removes the need for individual recapture and allows for the estimation of adult abundance using samples collected solely from juveniles, as well as samples obtained lethally through fishing or hunting (Bravington et al. 2016a, Hillary et al. 2018). While CKMR can theoretically leverage any relationship, the most common applications so far have focused on parent-offspring pairs (POPs) (Bravington et al. 2016b, Ruzzante et al. 2019, Marcy-Quay et al. 2020) and/or half-sibling pairs (HSPs) <u>(Hillary et al. 2018, Patterson et</u> <u>al. 2022a)</u>. Similar to conventional CMR, CKMR can estimate quantities beyond abundance, including survival (Hillary et al. 2018), fecundity (Bravington et al. 2016b), dispersal (Feutry et al. 2017, Conn et al. 2020, Patterson et al. 2022a), and, potentially, population growth rate, though which parameters can be estimated depends on the form of the model and type of kin pairs modeled. In cases where sampling is limited to juveniles, CKMR can provide added value to conventional CMR by generating parameter estimates for the adult population while CMR estimates parameters for the sampled (in this case juvenile) portion of the population. These advantages and possibilities make CKMR an exciting tool to improve monitoring efforts and population assessments for data-limited species of management and conservation concern, either in conjunction with, or in place of, conventional CMR.

Despite CKMR’s strong potential to provide key information for conservation and management, its implementation has been slowed by a lack of clarity regarding the flexibility and limitations of the method. Several studies have discussed factors that are likely to cause bias if left unaccounted for in CKMR models (Bravington et al. 2016a, Conn et al. 2020, Waples and Feutry 2021, Trenkel et al. 2022), but there have been few quantitative assessments of the biases that arise from applying an overly simplistic CKMR model to a population with complex dynamics (but see Conn et al. 2020, Waples and Feutry 2021). For example, a simple base-case CKMR model (e.g., Equations 3.3 and 3.10 in Bravington et al. 2016a) produces an abundance estimate that assumes abundance is constant over the modeled time period. However, real populations experience interannual fluctuations in population size. If such changes are persistent or severe (e.g., following an environmental disaster or introduction of heavy fishing pressure), then it will be necessary to specify a more complex CKMR model that can accommodate a changing population if year-specific abundance estimates are desired.

When modeling half-sibling relationships in particular, a simple base-case CKMR model assumes that the probability of two individuals sharing a parent is a simple exponential function of the year gap that separates their births. However, many long-lived species exhibit intermittent breeding whereby one or more years elapse between reproductive events (Morbey and Shuter 2013, Shaw and Levin 2013, Desprez et al. 2018, Bauwens and Claus 2019, Skjæraasen et al. 2020), resulting in different probabilities of detecting half-siblings depending on the age gap (Waples and Feutry 2021). Systematic intermittent breeding will cause bias in CKMR parameter estimates if unaccounted for in the model (Waples and Feutry 2021). While it may be possible to infer breeding periodicity based on the distribution of observed kin pairs in the data, instances of off-cycle breeding, mixed breeding schedules (e.g., a population comprising both annual and multiennial breeders), and aging uncertainty that leads to errors in cohort assignment may obscure the signal (Rivalan et al. 2005, Cubaynes et al. 2011, Öst et al. 2018, Higgs et al. 2020).

Finally, a core component of CKMR is the use of age data, which is required to assign individuals to the correct cohort (Bravington et al. 2016a). Direct aging is very challenging for some taxa (Cailliet 2015), and length-based age assignment is prone to bias when growth curves are based on size-selective sampling, as they often are (Gwinn et al. 2010). While more advanced statistical methods can account for uncertainty in aging during the modeling process (Schwarz and Runge 2009), it may also be possible to alleviate bias by targeting sampling to age classes that can be reliably aged, such as young-of-the-year (YOY) which are often easily distinguished from other age classes by their small size and/or the presence of umbilical scars (Feldheim et al. 2002). However, sampling constraints will not always permit long-term sampling of YOY and the number of cohorts required to produce robust parameter estimates with CKMR is unclear. A better understanding regarding the circumstances in which unobserved population dynamics or sampling limitations are likely to bias CKMR model estimates, in combination with strategies to mitigate that bias, will help ensure robust application of the method and facilitate its integration into conservation and management frameworks.

Elasmobranchs (sharks, skates, and rays) are a group of highly vulnerable marine species that play key ecological roles as apex- and meso-predators in ecosystems around the world (Vaudo and Heithaus 2011, Ferretti et al. 2018) and are likely to benefit from future application of CKMR. Around one-third of the 1200+ elasmobranch species are threatened with extinction, due primarily to overfishing (Dulvy et al. 2021), while nearly half of elasmobranch species (46%) are classified on the IUCN Red List of Threatened Species as Data Deficient and only a small fraction of exploited populations are managed sustainably (Kindsvater et al. 2018).

Conventional methods for estimating abundance and mortality are intractable for many elasmobranch populations because individual recapture rates for highly mobile elasmobranch species can be very low (Kohler and Turner 2001), and it can be logistically challenging to physically capture and mark larger species (Guttridge et al. 2017). In contrast to conventional methods, CKMR requires only small tissue samples that can be obtained from adults via biopsy, or from juveniles that are easier to handle than their adult counterparts. There is also no need for individual recapture so each animal only needs to be captured and handled once, making this a more feasible approach for many elasmobranch populations. In addition, when adults are unavailable for sampling, the life histories of many elasmobranch species allow for the use of juvenile-only CKMR models (e.g., half-sibling (HS) CKMR) that can estimate adult abundance without sampling a single adult <u>(Bravington et al. 2016a, Førland 2019)</u>. Considering that many migratory elasmobranchs use nursery areas where juveniles are more readily available for sampling than adults (Heupel et al. 2007), the potential for CKMR to provide novel insights into difficult-to-study elasmobranch populations is vast.

CKMR has been applied to several elasmobranch populations to date (Bradford et al. 2018, Hillary et al. 2018, Bravington et al. 2019, Delaval et al. 2022, Trenkel et al. 2022, Patterson et al. 2022a) and is likely to be an important tool to inform elasmobranch conservation and management in the future. However, elasmobranch populations are susceptible to steep population declines arising from overexploitation (Ferretti et al. 2018), commonly exhibit multiennial breeding cycles (Nosal et al. 2021), and are exceptionally challenging to age (Cailliet 2015). As such, there is a risk that CKMR models that do not sufficiently account for these factors will produce biased parameter estimates that will be unwittingly incorporated into management frameworks, leading to incorrect management actions that ultimately threaten elasmobranch populations.

To facilitate the robust application of CKMR to elasmobranchs and other long-lived taxa facing similar challenges with abundance estimation, we investigated the sensitivity of CKMR parameter estimates to unmodeled dynamics related to population growth and breeding schedule, as well as uncertainty in age assignment. We used stochastic individual-based simulation to generate distinct populations of lemon sharks (*Negaprion brevirostris*), a representative long-lived species with promiscuous mating and multiennial breeding, under different population dynamics scenarios and sampled each population using three sampling schemes that targeted different age classes. Two different CKMR models were fit to each dataset: one that was naïve to at least one component of the data-generating model (naïve model) and one that was adapted to account for all relevant population dynamics (adapted model). We compared the bias in parameter estimates from both models (naïve vs adapted) and across all three sampling schemes, including one in which age data were unreliable. Finally, we applied a model that was adapted for population growth and multiennial breeding to two decades of real genetic data from a small population of lemon sharks in Bimini, Bahamas, to generate a time-series of abundance estimates for the breeding population of females. Collectively, these results provide important insights into the ways in which unmodeled population dynamics, sampling selectivity, and aging error affect CKMR model performance, while also offering guidance regarding sampling design and model construction.

## 2. Methods

Our simulation framework comprised four primary components: 1) an individual-based population simulation that stochastically generated distinct populations with known parameters, 2) selective sampling of age classes from those populations, 3) construction of a pairwise comparison matrix from the samples, and 4) a CKMR model that was fit to the pairwise comparison matrix to estimate the known population parameters. The first three components comprised our data-generating model (DGM) while the latter formed our estimation model (Figure 1).

**Figure 1:**
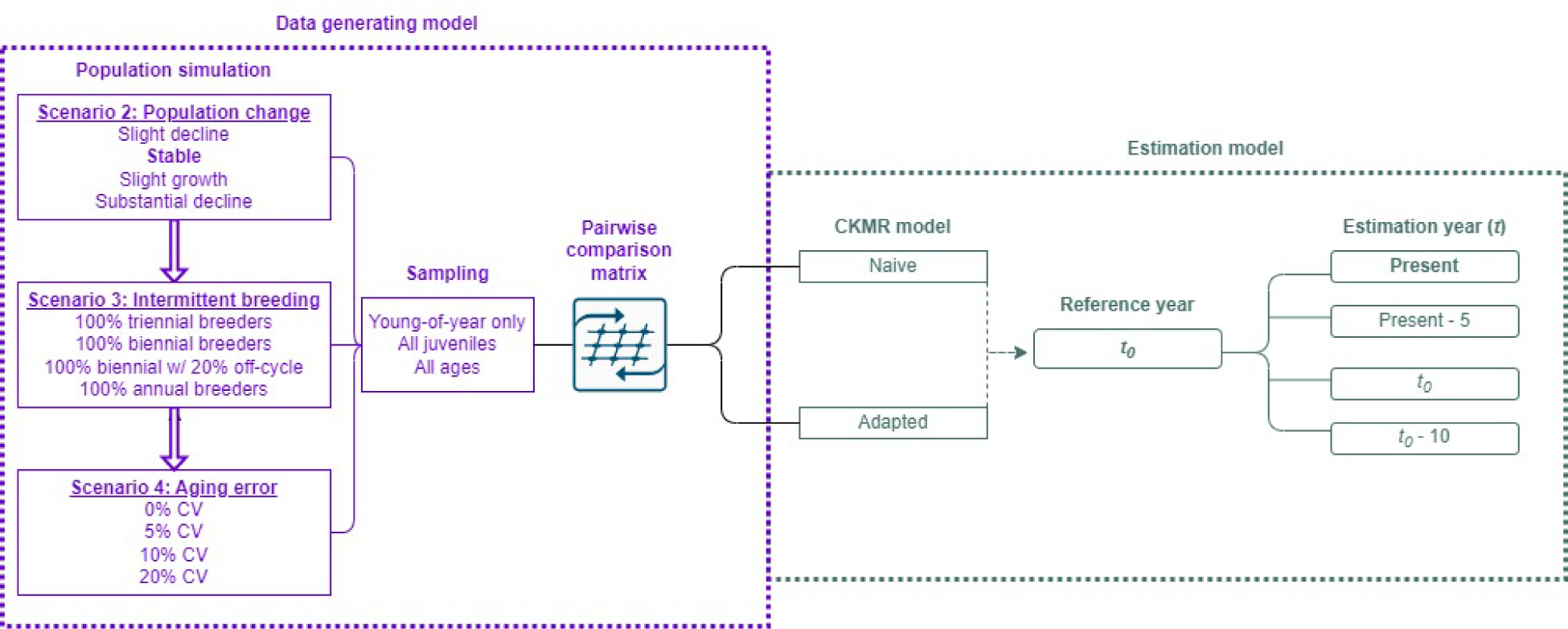
Schematic of CKMR sensitivity tests, examined via individual-based simulation (see also Tables 1 and 2; Scenario 1 was model validation, and Scenario 5 involved real genetic data, so are not included here). Populations with distinct pedigrees were produced and sampled via an individual-based data-generating model (purple). Population parameters were individually varied for each of three scenarios. Each population was sampled in three ways, and each set of samples was used as input to two estimation models (green): one model was naïve to the added population dynamics of the DGM, and one model was adapted to account for them. The year of estimation (year *t*) was varied for Scenario 2; otherwise, simulation results that are discussed in the text used the model settings highlighted in bold.

We then tested the interplay of population dynamics and model complexity by iteratively varying a subset of population parameters (Table 1) and fitting two CKMR models to the data: one that was naïve to the added dynamics, and one that was adapted to account for them. Each scenario was repeated 500 times, with each iteration producing a population with a distinct pedigree and parameter estimates.

**Table 1:** Simulation scenarios for data-generating model and estimation model.

| Scenario | Test | 10<br>year<br>mean<br>$\lambda$ | Model<br>equation(s)<br>used | Parameters<br>estimated | Parameters<br>derived | Parameters<br>compared<br>(naïve vs<br>adapted) |
| --- | --- | --- | --- | --- | --- | --- |
| 1.1 | Model validation | 1 | Eq. 2 | $N_{\varphi}, N_{\sigma}, \square$ | NA | $N_{\varphi}, N_{\sigma}, \square$ |
| 1.2 | Misidentify<br>aunt/niece pairs as<br>HSPs | 1 | Eq. 2 | $N_{\varphi}, N_{\sigma}, \square$ | NA | $N_{\varphi}, N_{\sigma}, \square$ |
| 2.1 | Slight population<br>decline | 0.99 | Eq. 2, Eq. 3,<br>Eq. 7, Eq. 8 | $N_{\varphi(t0)}, N_{\sigma(t0)}, \square, \lambda$ | $N_{\varphi(t)}, N_{\sigma(t)}$ | $N_{\varphi(t)}, \square, \lambda$ |
| 2.2 | Slight population<br>growth | 1.01 | Eq. 2, Eq. 3,<br>Eq. 7, Eq. 8 | $N_{\varphi(t0)}, N_{\sigma(t0)}, \square, \lambda$ | $N_{\varphi(t)}, N_{\sigma(t)}$ | $N_{\varphi(t)}, \square, \lambda$ |
| 2.3 | Severe population<br>decline | 0.93 | Eq. 2, Eq. 3,<br>Eq. 7, Eq. 8 | $N_{\varphi(t0)}, N_{\sigma(t0)}, \square, \lambda$ | $N_{\varphi(t)}, N_{\sigma(t)}$ | $N_{\varphi(t)}, \square, \lambda$ |
| 2.4 | Stable population<br>growth | 1 | Eq. 2, Eq. 3,<br>Eq. 7, Eq. 8 | $N_{\varphi(t0)}, N_{\sigma(t0)}, \square, \lambda$ | $N_{\varphi(t)}, N_{\sigma(t)}$ | $N_{\varphi(t)}, \square, \lambda$ |
| 3.1 | 100% biennial<br>breeders | 1 | Eq. 3, Eq. 4,<br>Eq. 7, Eq. 8 | $N_{\varphi(t0)}, N_{\sigma(t0)}, \square, \lambda,$<br>$\psi$ | $N_{\varphi(t)}, N_{\sigma(t)}$ | $N_{\varphi(t)}, N_{\sigma(t)},$<br>$\square, \psi$ |
| 3.2 | 100% biennial<br>breeders w/<br>stochastic off-cycle breeding | 1 | Eq. 3, Eq. 4,<br>Eq. 7, Eq. 8 | $N_{\varphi(t0)}, N_{\sigma(t0)}, \square, \lambda,$<br>$\psi$ | $N_{\varphi(t)}, N_{\sigma(t)}$ | $N_{\varphi(t)}, N_{\sigma(t)},$<br>$\square, \psi$ |
| 3.3 | 100% triennial<br>breeders | 1 | Eq. 3, Eq. 4,<br>Eq. 7, Eq. 8 | $N_{\varphi(t0)}, N_{\sigma(t0)}, \square, \lambda,$<br>$\psi$ | $N_{\varphi(t)}, N_{\sigma(t)}$ | $N_{\varphi(t)}, N_{\sigma(t)},$<br>$\square, \psi$ |
| 3.4 | 100% annual<br>breeders | 1 | Eq. 3, Eq. 4,<br>Eq. 7, Eq. 8 | $N_{\varphi(t0)}, N_{\sigma(t0)}, \square, \lambda,$<br>$\psi$ | $N_{\varphi(t)}, N_{\sigma(t)}$ | $N_{\varphi(t)}, N_{\sigma(t)},$<br>$\square, \psi$ |
| 4.1 | Minimal age-length uncertainty<br>(5% CV) | 1 | Eq. 3, Eq. 4,<br>Eq. 7, Eq. 8 | $N_{\varphi(0)(t0)}, N_{\sigma(0)(t0)},$<br>$\square, \lambda, \psi$ | $N_{\varphi(t)}, N_{\sigma(t)}$ | $N_{\varphi(t)}, \square$ |
| 4.2 | Moderate age-length uncertainty<br>(10% CV) | 1 | Eq. 3, Eq. 4,<br>Eq. 7, Eq. 8 | $N_{\varphi(0)(t0)}, N_{\sigma(0)(t0)},$<br>$\square, \lambda, \psi$ | $N_{\varphi(t)}, N_{\sigma(t)}$ | $N_{\varphi(t)}, \square$ |
| 4.3 | Substantial age-length uncertainty<br>(20% CV) | 1 | Eq. 3, Eq. 4,<br>Eq. 7, Eq. 8 | $N_{\varphi(0)(t0)}, N_{\sigma(0)(t0)},$<br>$\square, \lambda, \psi$ | $N_{\varphi(t)}, N_{\sigma(t)}$ | $N_{\varphi(t)}, \square$ |

*Note:* See Table 2 for parameter definitions.

### 2.1 Data generating model

Parameters governing our individual-based population simulations were designed to replicate the life history traits and population dynamics of lemon sharks in Bimini, Bahamas, <u>(similar to White et al. 2014</u>) (Appendix S1: Table S1). Females bred with one, two, or three distinct males each breeding cycle and produced two or three pups with each male, resulting in a range of 2-9 total pups produced per female per year. We set no limit on the number of females an individual male could breed with. As a consequence, the variance in reproductive output for males was much greater, ranging from 2 – 41 offspring per breeding male per year (median = 6). After maturity, fecundity was age-invariant, so sex-specific lifetime reproductive output was approximately equal across the population. Survival was assumed constant within each of three life stages, which we designated as young-of-year (YOY; age 0), juvenile (age 1-11) and adult (age 12-50). We assigned knife-edged maturity following White et al. (2014), so every individual age 12 and over was available for breeding, while no individuals younger than age 12 were allowed to breed.

#### 2.1.1 Population growth

We varied population growth in our DGM by reducing or adding mortality on juveniles and adults (Appendix S1: Table S1). For the slight increase and decline scenarios (Table 1: scenarios 2.1 and 2.2), mortality was increased or reduced by ∼1% from the beginning of the simulation, resulting in a population growth rate of +/- 1% per year. To achieve more substantial declines in population size (Table 1: scenario 2.3), we simulated a stable population for 80 years and then stochastically imposed 4-7% added mortality for juvenile and adult age classes for years 81-90 (when imposed from the beginning of the simulation, the population invariably went extinct). This produced a population that declined at a rate of ∼7% per year for the last 10 years of the simulation.

#### 2.1.2 Intermittent breeding

Many elasmobranchs systematically breed on multiennial cycles (Feldheim et al. 2002, 2017, Nosal et al. 2021). To examine the bias that accrues when this trait is unaccounted for in a CKMR model, we ran simulations where all females bred on a bi- or triennial cycle (Table 1: scenario 3.1 – 3.3), including one scenario where we allowed 20% of breeding females to stochastically breed off-cycle or fail to breed when they were on-cycle (scenario 3.2). Each female in our simulation was assigned a breeding cycle at birth, which determined the first year of reproduction for multiennial breeders. In scenarios with biennial breeders, this resulted in a population where half of the females reproduced for the first time in the year they matured (age 12) and the other half reproduced for the first time the following year (age 13). For the scenario with triennial breeding, a third group reproduced for the first time at age 14. Mature males were assumed available to breed every year once they reached maturity at age 12.

### 2.2 Sampling

All simulated populations were sampled using three different schemes that selected for different age classes: the first drew samples exclusively from young-of-year (age 0) individuals; the second made juveniles of all ages except young-of-year (ages 1-11) available to sample; and the third allowed sampling of all age classes (ages 0-50). These scenarios were chosen to replicate potential sampling opportunities for elasmobranchs such as nursery areas (Feldheim et al. 2002, Heupel et al. 2007), juvenile aggregation sites (Rowat et al. 2007, Jacoby et al. 2012), and resident populations (Snelson and Williams 1981), respectively.

In each case, the population was initially sampled at four different intensities representing 0.5%, 1%, 1.5%, and 2% of the population. Samples were drawn annually and non-lethally for four years at the end of the population simulation (i.e., years 87 – 90), following reproduction but before mortality each year. With a stable population, sampling 1.5% of the population resulted in an average of 616 total samples and 100-200 half-sibling pairs (HSPs), which is expected to produce a reasonable CV for all three sampling schemes (Bravington et al. 2016a). Therefore, following model validation, we focused on sampling 1.5% of the population for the remainder of our simulations.

#### 2.2.1 Aging uncertainty

A crucial component of CKMR is accurate aging, yet some taxa, including elasmobranchs, are notoriously difficult to age, with most efforts relying on length-at-age growth curves to assign age to sampled individuals (Cailliet 2015). To examine how imprecision in growth curves affects CKMR parameter estimates, we first constructed an age-length key for lemon sharks using data from a long-term study of the population in Bimini, Bahamas (Feldheim et al. 2014), and calculated the standard deviation of lengths for individuals with known ages, the majority of which (>95%) spanned ages 0-3. We then simulated lengths for each sampled individual (which were assigned ages in our DGM) using a von Bertalanffy growth curve for the species (Brown and Gruber 1988). Each individual was assigned a length by drawing a value from a normal distribution with the mean length-at-age specified by the von Bertalanffy curve, and the standard deviation derived empirically from our age-length key for individuals aged 0-2, and arbitrarily from a CV of 5%, 10%, or 20% for individuals aged 3+. After assigning lengths to each individual, we used a reverse von Bertalanffy growth curve with the same values for theoretical age of zero size (*t_0_* = −2.302), asymptotic average length (*L_inf_* = 317.65), and the growth coefficient (*K* = 0.057) and then re-assigned ages to sampled individuals based on their lengths, rounding to the nearest integer. This produced plausible, yet sometimes incorrect, ages (similar to age-slicing; see Ailloud et al. 2015, e.g.). The re-assigned ages were then used to construct the pairwise comparison matrix that was input to the CKMR model.

### 2.3 Pairwise comparison matrix

CKMR produces estimates of abundance and other population parameters by defining kinship probabilities for every pair of sampled individuals given relevant covariates (e.g., age, sex). We constructed two standard pairwise comparison matrices for each set of samples. The first matrix contained positive and negative kinship assignments for half-siblings. To satisfy the assumption of independent sampling, whenever full siblings or self-recaptures were present, all but one individual/instance was removed prior to construction of the matrix. Once the matrix was created, within-cohort comparisons were removed. Though CKMR models can be adapted to incorporate within-cohort comparisons (Førland 2019), without considerable modifications to the equations, within-cohort and cross-cohort comparisons will estimate different quantities (Waples and Feutry 2021). As such, removing within-cohort comparisons is common practice for application of HS CKMR <u>(Bravington et al. 2016a</u>). Kinship assignment in our simulations was known without error, so each comparison was assigned as a positive if the two individuals being compared were a half-sibling pair, and negative if not. Because half-siblings are genetically indistinguishable from aunt/niece (uncle/nephew, etc.) pairs, we included one scenario (Table 1: scenario 1.2) where we allowed such comparisons to contaminate the pool of half-siblings and evaluated the degree to which these false positives affected parameter estimates.

The second matrix was composed of parent-offspring (PO) comparisons, which was only relevant to the scenario that included sampling of adults. For each birth year represented in the dataset, individuals that were alive in that year were split into potential offspring or parents based on whether they were born in that year (potential offspring), reproductively mature at the time (potential parent), or neither, in which case they were left out of the matrix corresponding to that year. Each comparison was assigned as a positive if they were related as parent-offspring or negative if not.

Once the appropriate half-sibling and parent-offspring comparisons were defined, all matrices were collated and grouped by 1) type of relationship (half-sibling or parent-offspring), 2) birth year of younger individual in each comparison (a.k.a. *y_j_*; see Section 2.4.1), 3) reference year gap (a.k.a. (*y_j_ – t_0_*); see Section 2.4.2), and 4) birth year gap (a.k.a. δ; see Section 2.4.1), as applicable. The number of observed kin pairs (*Y*) was then modeled as a random variable, with the probability of success defined by Equations 1-9 below, and *n* equal to the total number of comparisons in each group (see Appendix S1: S1.1 and Table S2 for more details on the pairwise comparison matrix, and Appendix S1: S1.2 for more details on kinship types that can cause issues for half-sibling CKMR e.g., aunt/niece pairs).

### 2.4 Estimation models

Kinship probabilities for each pairwise comparison in CKMR are derived from the expected reproductive output of individual animals (defined by covariates such as age and sex) relative to the total reproductive output of the population in the birth year of the younger individual in each pairwise comparison (Bravington et al. 2016a). The specific equations we used to define kinship probabilities in our CKMR models varied with the scenarios we tested, with each scenario comparing a “naïve” model to an “adapted” model, where the naïve model ignored one key dynamic of the simulated population and the adapted model accounted for it. Our equations are based on the general equations defined in Bravington et al. (2016a).

#### 2.4.1 Base-case CKMR model

Let *P{K_i,j_ = MHSP}* be the probability that individuals *i* and *j* are a maternal half-sibling pair (i.e. they share a mother but not a father). Probabilities for *K_i,j_* depend on the likelihood that the same individual that birthed the older offspring (*i*) survived and gave birth to the younger offspring (*j*). If we assume that all animals of reproductive age in the population during *i* and *j*’s birth years are equally likely to have birthed each of them, then the probability of kinship (*K*) can be defined as

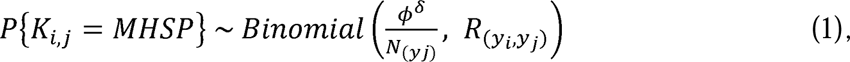

where,

*ϕ* is the annual survival probability for adults,

*δ* is the number of years between the birth years of individuals *i* and *j* (i.e. *y_j_* – *y_i_*) during which any potential parent of *i* may have died a.k.a. the “birth year gap”,

*y*_j_ is the birth year of individual *j* (the younger sibling),

*R*_(*yi,yj*)_ reflects the total number of pairwise comparisons between individuals born in years *y_i_* and *y_j_*, and

*N* _(*yj)*_ is the total number of mature females in year *y_j_*.

Now, let *P{K_i,j_ = MPOP}* refer to the probability that individuals *i* and *j* are related as a maternal parent-offspring pair (MPOP). In this case, survival only enters the equation for MPOPs if sampling is non-lethal (as it was in our simulations) *and* if the potential parent was sampled before the potential offspring was born. If, on the other hand, the potential mother *i* was captured in or after offspring *j*’s birth year and was reproductively mature, then we know that she was alive in the year the offspring was born and, assuming constant fecundity across the population, is equally likely to have birthed *j* as any other potential mother. Assuming the pool of potential parents was filtered to only include individuals that were mature and were not known to have died before *j*’s birth year, the probability of kinship is

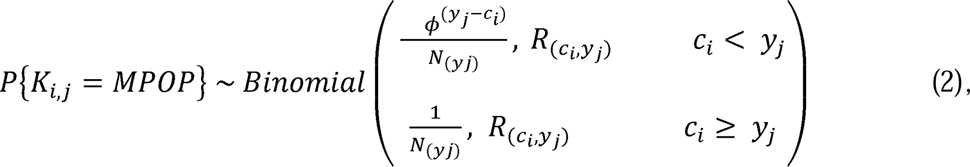

where *c_i_* refers to the year in which the potential parent was captured. For each offspring birth year (*y_j_*), it is crucial to ensure that individuals who were not reproductively mature are not included as potential parents for that year. This restriction can be added directly to the model <u>(Bravington et al. 2016a, Conn et al. 2020)</u> or implemented during construction of the pairwise comparison matrix, as was done here.

Given that sampling was nonlethal for the parent, then mother *i*’s survival to the year of *j*’s birth is conflated with detection probability (when *c_i_ < y_j_*). In circumstances where the individual recapture rate of adults is non-negligible, an additional parameter defining the adult detection probability will be required to disentangle the state (□) and observation (detection probability) processes. However, if sampling is sparse such that individual recaptures are exceedingly rare, the cost of estimating an extra parameter for detection probability likely outweighs the benefits.

Equations 1 and 2 define our base-case CKMR model. Though the probabilities presented here focus on maternal kinship, the same probabilities apply to males and paternal kinship (see Equations 8 and 9 below). While all parameter estimates presented here incorporate HSPs, POPs were only included in the likelihood for the sampling scheme in which adults were sampled with all other age classes; otherwise, the likelihood included HS kinship probabilities only.

#### 2.4.2 Population growth model

To account for population growth/decline in our CKMR model, we defined a simple exponential growth model to describe the population dynamics, where

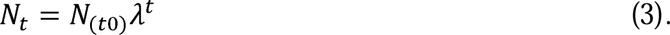

As such, the kinship probabilities become:

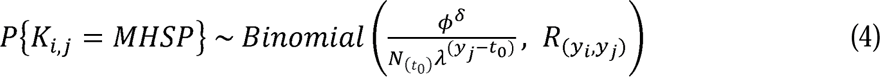

and

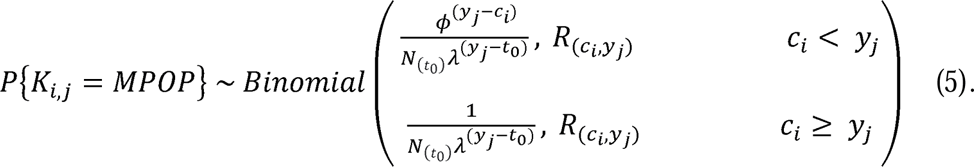

Here, *λ* defines the annual population growth rate and *t_0_*refers to the initial model year, also called the “reference year” <u>(Bravington et al. 2016a)</u>. The reference year typically refers to the earliest instance of *y_j_* in the pairwise comparison matrix but could refer to any modeled year. To assess the capacity of CKMR to generate year-specific abundance estimates, we fit the same CKMR model to each dataset four times. In each instance, we set the reference year (*t_0_*) to the earliest instance of *y_j_*. Then, we estimated *N_t_*10 years prior to the reference year (before data were collected), in the reference year (*t_0_*), 5 years prior to the last year of the simulation, and the last year of the simulation (i.e., present). We also tested two different methods for generating year-specific abundance estimates: one where *t_0_* was fixed to the first year of data (first instance of *y_j_*) and *N_t_*was calculated as a derived quantity (Table 2 – our primary approach), and one where *t_0_* was directly set to the year of interest (i.e., *N* was directly estimated in year *t*; see Appendix S1: S1.3 for more discussion on CKMR with a changing population).

**Table 2:** Model parameters and priors.

| Parameter | Definition | Prior |
| --- | --- | --- |
| $N_{s(t0)}$ | Sex-specific abundance in year 0. | $N_{s(t0)} \sim \text{Normal}(\mu, \sigma)$<br>$\mu \sim \text{Uniform}(1, 10000)$<br>$\sigma \sim \text{Uniform}(1, 10000)$ |
| $\phi$ | Annual survival | Uniform(0.5, 0.95): default<br>Uniform(0.5, 0.99): for small population simulations and application to Bimini data |
| $\lambda$ | Annual finite population growth rate | Uniform(0.95, 1.05): default<br>Uniform(0.80, 1.20): for severe decline scenario<br>Uniform(0.70, 1.30): for small population simulations and application to Bimini data |
| $\psi$ | Proportion of individuals that breed every $a$ years | Uniform(0, 1) |
| $a$ | Years between breeding | Fixed |
| **Derived quantities** | **Definition** | **Equation** |
| $N_{s(t)}$ | Sex-specific abundance in year $t$ | $N_{s(t0)} * \lambda^{(yt - t0)}$ |
| $N_{\varphi b(t)}$ | Abundance of breeding females in year $t$ (in multiennial simulations) | $\frac{a + \psi - a\psi}{a} N_{(t)}$ |

*Note:* $N_s$ is a general term that encompasses both $N_{\varphi}$ , $N_{\sigma}$ when the sex-specific parameters were treated the same.

#### 2.4.3 Intermittent breeding model

If a population – or subset of a population – systematically breeds on a non-annual schedule, then CKMR estimates will be biased unless this behavior is accounted for in the model (Waples and Feutry 2021). We accounted for intermittent breeding dynamics in our CKMR model via the inclusion of parameters *a* and Ψ, where *a* refers to the number of years between breeding (e.g., 2 for biennial breeders), and Ψ is the proportion of individuals that breed every *a* years (similar to <u>Patterson et al. 2022b)</u>. This implies that (*1* - Ψ) individuals breed annually. We assume that the proportion of on-cycle breeders that breed in a given year is *1/a*. Thus, the effective number of female breeders in a given year (*Ñ_t_*) is given by

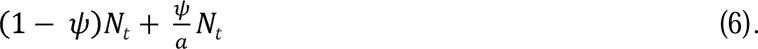

Accounting for interannual population dynamics (Eq. 3), the full probability of maternal half-sibling kinship for a population that reproduces on a multiennial schedule becomes

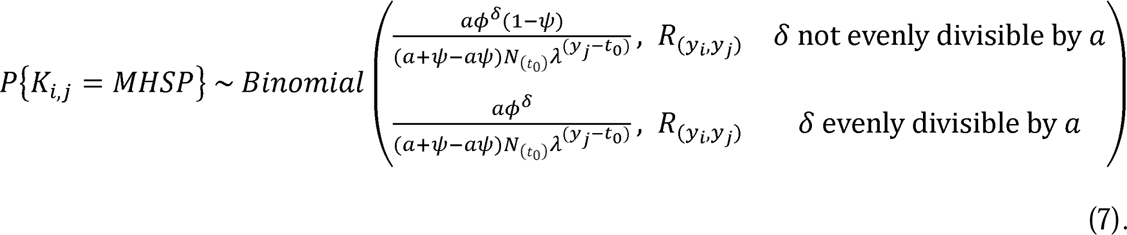

If 100% of females breed on a biennial cycle (i.e. *a* = 2 and *Ψ* = 1), then the probability of finding half-siblings that are separated by an odd number of birth years is 0. It is the presence of δ intervals that are not evenly divisible by *a* that provide information on the parameter *Ψ* (see Appendix S1: S1.4). Alternatively, when a consistent pattern is observed in the year gaps that separate HSPs, one may be tempted to remove off-cycle comparisons from the pairwise comparison matrix - since they have no chance of revealing a positive comparison - and fit a model that is naïve to intermittent breeding (e.g., Eq. 4) to the on-cycle comparisons only. We thus evaluated two different types of “naïve” model: one where we retained off-cycle comparisons in the pairwise comparison matrix (the “naïve” model), and one where we removed off-cycle comparisons (the “naïve - filtered” model).

In our simulations, intermittent breeding dynamics were only present for females, and all males in the population were available for breeding each year; as such, Equation 7 only applied to maternal comparisons, while kinship probabilities for paternal half sibling pairs (PHSPs) continued to be defined as:

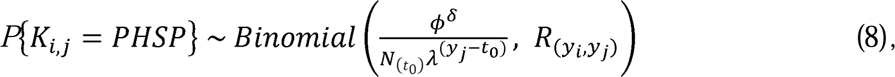

which is the same as the maternal kinship probability defined in Equation 4.

Because the parent is directly sampled in PO CKMR, there is no need to explicitly account for breeding periodicity in the likelihood; therefore, we continued to use Equation 5 for maternal PO comparisons when applicable. Similarly, kinship probabilities for paternal parent-offspring pairs (PPOPs) mirrored those for MPOPs in Equation 5:

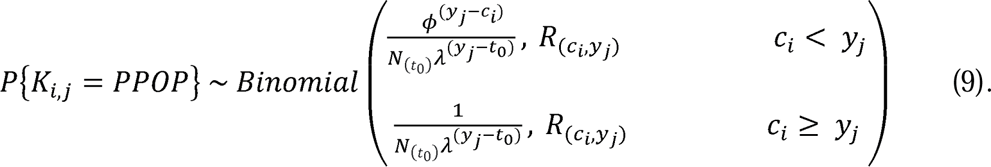

For all multiennial simulations, we simulated a population with an approximately stable growth rate, set the reference year (*t_0_*) to the earliest instance of *y_j_*, and estimated *N_t_*in the present (i.e., most recent year of sampling), as we expect this to be a common approach in real-life applications of CKMR.

#### 2.4.4 Estimation framework

We adopted a Bayesian approach to CKMR parameter estimation, which allows for the incorporation of auxiliary data and/or expert knowledge as priors on model parameters (Kéry and Schaub, 2012b). For the scenarios tested here, survival and other parameters were assigned reasonably diffuse priors to reflect data-limited situations (Table 2). Though it is possible to estimate sex-specific survival (□) and population growth rates (λ), these parameters were shared between males and females in our models. The posterior distributions for parameters were approximated using Markov Chain Monte Carlo (MCMC) sampling, implemented using the software JAGS (Plummer 2003) and applied in the *R* environment (Denwood 2016, R Core Team 2021). We ran two Markov chains with a thinning rate of 20, drawing 40,000 samples from the posterior distribution following a burn-in of 50,000 samples. These settings were empirically derived by assessing autocorrelation among successive draws and convergence among the chains. We assessed convergence of the final Markov chains with trace plots and the Gelman-Rubin statistic (Gelman and Rubin 1992), and removed from further analysis any simulation replicate with an Rhat value > 1.01, although these instances were rare (∼1.5% of simulations).

### 2.5 Application to lemon sharks

#### 2.5.1 Bimini lemon shark dataset

A long-term genetic dataset from lemon sharks in Bimini, Bahamas, was used to illustrate application of our multiennial CKMR model (Eq. 7) to a dataset derived entirely from juvenile tissue samples (Feldheim et al. 2014). Lemon sharks are large viviparous (live-bearing) elasmobranchs that reach sexual maturity at approximately 12 years of age (Brown and Gruber 1988) with a lifespan exceeding 30 years (Brooks et al. 2016). Female lemon sharks at Bimini are regionally philopatric and return to Bimini to pup on a biennial schedule, while the males with which they mate likely reproduce over a much larger area (Feldheim et al. 2002). Juveniles use the shallow waters surrounding Bimini as a nursery and remain in the area until 2-3 years of age or until they reach 90 cm in length (Morrissey and Gruber 1993) and generally do not move between the North and South Islands (Gruber et al. 2001). The Bimini nursery contributes to a larger Western Atlantic population that is classified as Vulnerable on the IUCN Red List (Hansell et al. 2018, 2021, Carlson et al. 2021). The Bimini nursery has been intensively studied since 1995, with an estimated 99% of newborn sharks sampled at the Bimini North Island each year (Dibattista et al. 2007). The ability to heavily sample multiple litters has allowed for reliable reconstruction of maternal genotypes, while paternal genotype reconstruction has often relied on relatively few newborns, resulting in high confidence in maternal kinship assignment and lower confidence in paternal kinship (Feldheim et al. 2002, 2004, 2014).

Given the disparities in kinship assignment and breeding range, we focused our CKMR model on maternal comparisons to estimate abundance and survival of adult females. We used samples collected from the North Island, a small isolated nursery for lemon sharks aged 0-3 years old (Chapman et al. 2009), from 1993-2015. Most individuals in our dataset were sampled as YOY (92%) and easily identified by the presence of umbilical scars, so their ages were known. We estimated abundance of total females in our CKMR model using Equation 7, and derived the number of effective female breeders in each year using Equation 6. Thus, our scope of inference for parameter estimation encompassed the adult females that visited the North Island nursery to give birth during each modeled year, a number which is likely very small (White et al. 2014). We excluded sampled individuals without a known birth year from analysis as well as same-cohort comparisons (Bravington et al. 2016a), and any individuals for which maternal kinship assignment was uncertain.

There were many full sibling pairs in the dataset (1,515 individuals contributing to 1,129 pairs), but very few cross-cohort full siblings (only 4% of full sibling pairs). Including more than one individual from each litter in a CKMR analysis can result in non-independence among pairwise comparisons and unreliable estimates of variance, especially in small populations where sampling effort may be high relative to the population size <u>(i.e., non-sparse sampling; Bravington</u> <u>et al. 2016a, 2019)</u>. Because Bimini lemon sharks were exhaustively sampled with relatively few instances of cross-cohort full siblings, we hypothesized that the retention of littermates might provide valuable data to the CKMR model, even if it reduced the reliability of variance estimates. Therefore, we fit our multiennial CKMR model (Eq. 7) to two sets of data: one where we included full littermates in the analysis (though we still removed all within-cohort comparisons from the pairwise comparison matrix) and one where we only retained one individual per mother/sire breeding pair, similar to our approach with the larger simulated populations.

Finally, recognizing that the Bimini lemon shark dataset is unique in how thoroughly the population was sampled, we also examined whether the model performed similarly with a sparser dataset by randomly downsampling and reducing the number of samples from each year to 30% of the full dataset. To account for random variation surrounding which samples were retained, we iterated over the process 50 times, fit a CKMR model to each set of samples, and report the average of the median and 95% highest posterior density intervals (HPDI) of the 50 posterior distributions.

#### 2.5.2 Bimini lemon shark simulations

Preliminary application of our multiennial CKMR model (Eq. 7) to the real Bimini dataset suggested the population likely experienced alternating periods of growth and decline during the modeled period. Our demographic model (Eq. 3) assumes the population is growing exponentially, and we suspected that this may result in an averaging effect and imprecise parameter estimates over our multi-decadal time series of data, especially if an inconsistent trend was present. To test this hypothesis and further examine the effects of applying our model to a small population, we refined our DGM to produce a population of similar size and with similar dynamics as Bimini lemon sharks (Feldheim et al. 2002, DiBattista et al. 2011, White et al. 2014) and then sampled 90% of the YOY from that population over 20 years to achieve a dataset that resembled the real dataset.

We fit the first CKMR model after four years of sampling. Then, to replicate the type of real-time estimates that could be produced by integrating CKMR into long-term monitoring efforts, we iteratively added one year of samples to the dataset until reaching the end of the time series, fitting three CKMR models each time a year of samples was added: one that included all samples that had been collected up to the most recent year of sampling, one that subset for samples within a five-year window of the most recent year of sampling, and one that subset for samples within a three-year window. In each case, the reference year (*t_0_*) was set to the birth year of the second oldest individual in the dataset being used (i.e., the first instance of *y_j_*), and abundance (*N*_♀*t*_) was estimated in the most recent year of sampling following Eq. 3. Abundance trend was then tracked in two ways: via a time-series of female abundance, and estimation of λ over the window of included samples. Finally, we applied the same approach to the real genetic dataset to estimate the total number of adult females, then derived the number of year-specific female breeders using Equation 6, and compared our estimates of yearly female breeders to estimates that have been independently obtained for the population using a reconstructed pedigree (DiBattista et al. 2011).

## 3. Results

### 3.1 Model validation

When the assumptions of the base-case model (Eqs. 1 and 2) were met, CKMR produced unbiased estimates of abundance under all sampling schemes and intensities (Figure 2a), with increasing precision as sampling intensity increased (Figure 2b). The model produced unbiased estimates of abundance whether the likelihood included HSPs only (as in the “sample juveniles” and “target YOY” scenarios) or jointly considered HSPs and POPs (as in the “sample all ages” scenario), though we note improved precision for the latter. At very low sampling intensities (0.5% of the target population sampled), fewer than 25 HSPs were identified for all sampling schemes (Figure 2c) and fewer than 5 parent-offspring pairs (POPs) were identified for the sampling scheme that included all ages (Figure 2d). In contrast, when 2% of the population was sampled, over 200 HSPs were identified on average for all sampling schemes, while 10-40 POPs were identified for the scenario in which all age classes were sampled. Including aunt/niece (uncle/nephew, etc.) pairs as HSPs had a minimal effect on abundance estimation, as such instances were rare in our dataset (Appendix S1: S1.2, Figure S1); similarly, while same-cohort full siblings were common, instances of cross-cohort full siblings were rare (Appendix S1, Figure S2). These results demonstrate that a simple base-case CKMR model can produce unbiased abundance estimates across a range of potential sampling scenarios when population dynamics align with the model’s assumptions while suggesting that false positive HSPs arising from misidentified aunt/niece (uncle/nephew, etc.) pairs are likely to be rare for randomly sampled long-lived promiscuous species.

**Figure 2:**
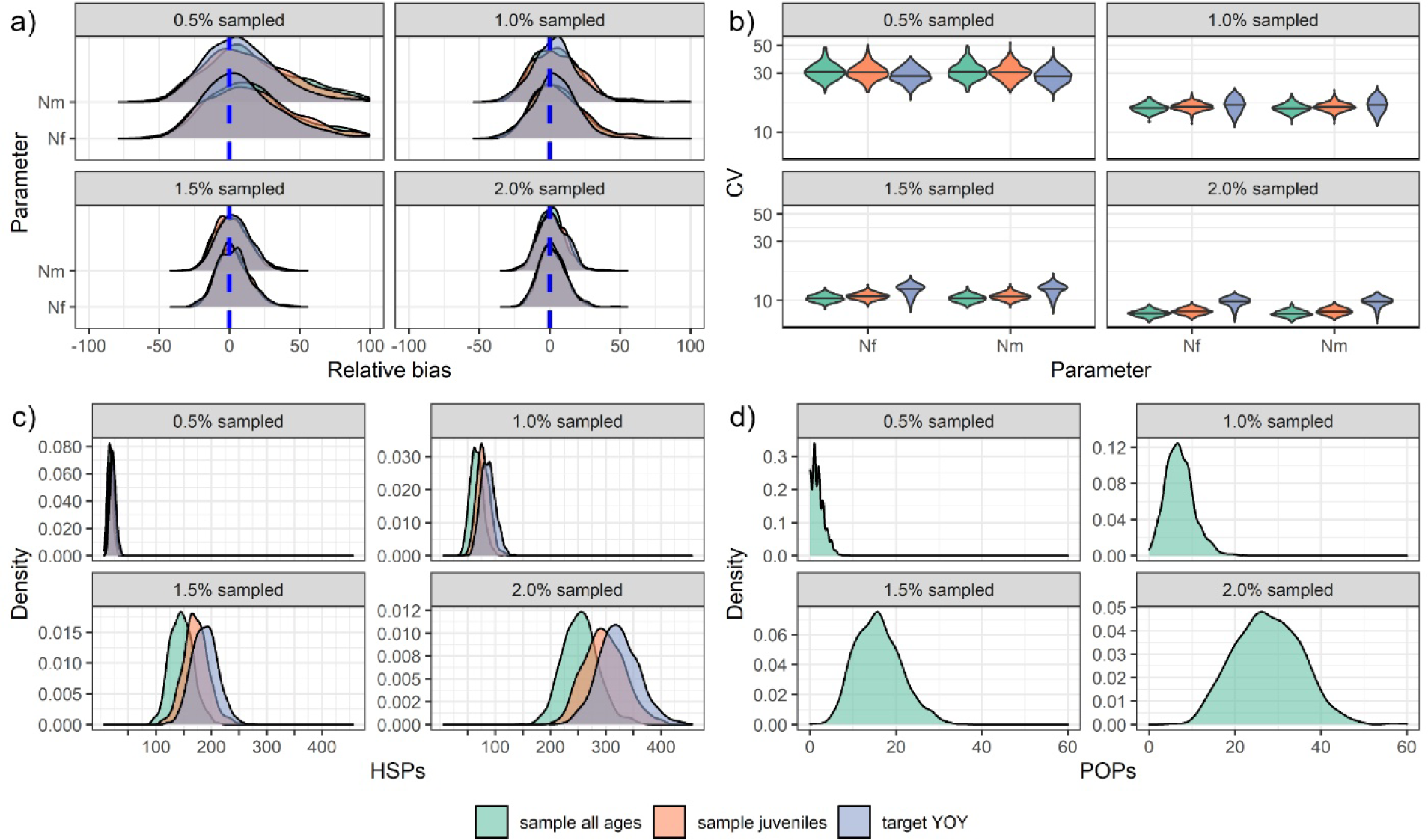
Base CKMR model performance and kin pairs detected for three different sampling schemes at four different sampling intensities over 500 iterations. **a)** Relative bias of abundance estimates of adult females (*Nf*) and males (*Nm*) as a percentage of the truth (i.e. relative bias x 100). Bias was calculated from the median of each of 500 posterior distributions. **b)** CV on abundance estimates with log-scaled Y axis for visualization. **c)** Number of sex-specific half-sibling pairs detected by sampling scheme and sampling intensity. For each iteration, the number of half sibling pairs for each sex was calculated and averaged. **d)** Number of sex-specific parent-offspring pairs detected for the “sample all ages” sampling scheme.

### 3.2 Population growth

When we simulated a population with a trend that was growing or declining in size and compared year-specific truths to the abundance estimates generated by our base-case CKMR model that was naïve to a population trend (Figure 3a-c, orange), the disparity between the quantities grew the further the estimation year (year *t*) was projected into the past. These disparities were rectified by adapting our CKMR model to include a population growth model (Eq. 3) and deriving *N_t_* from estimates of *N_(t0)_* and λ (Figure 3a-c, blue; Table 2). Uncertainty accrued as *N_t_*was projected further from the mode of the data (Figure 3d), even with a stable population (Appendix S1, figure S3a). When we varied *N_(t0)_* to generate year-specific abundance estimates rather than deriving *N_t_*, the model showed similar, though not identical, trends (Appendix S1: S1.3, figure S4), suggesting that the two approaches are functionally equivalent in most cases.

**Figure 3:**
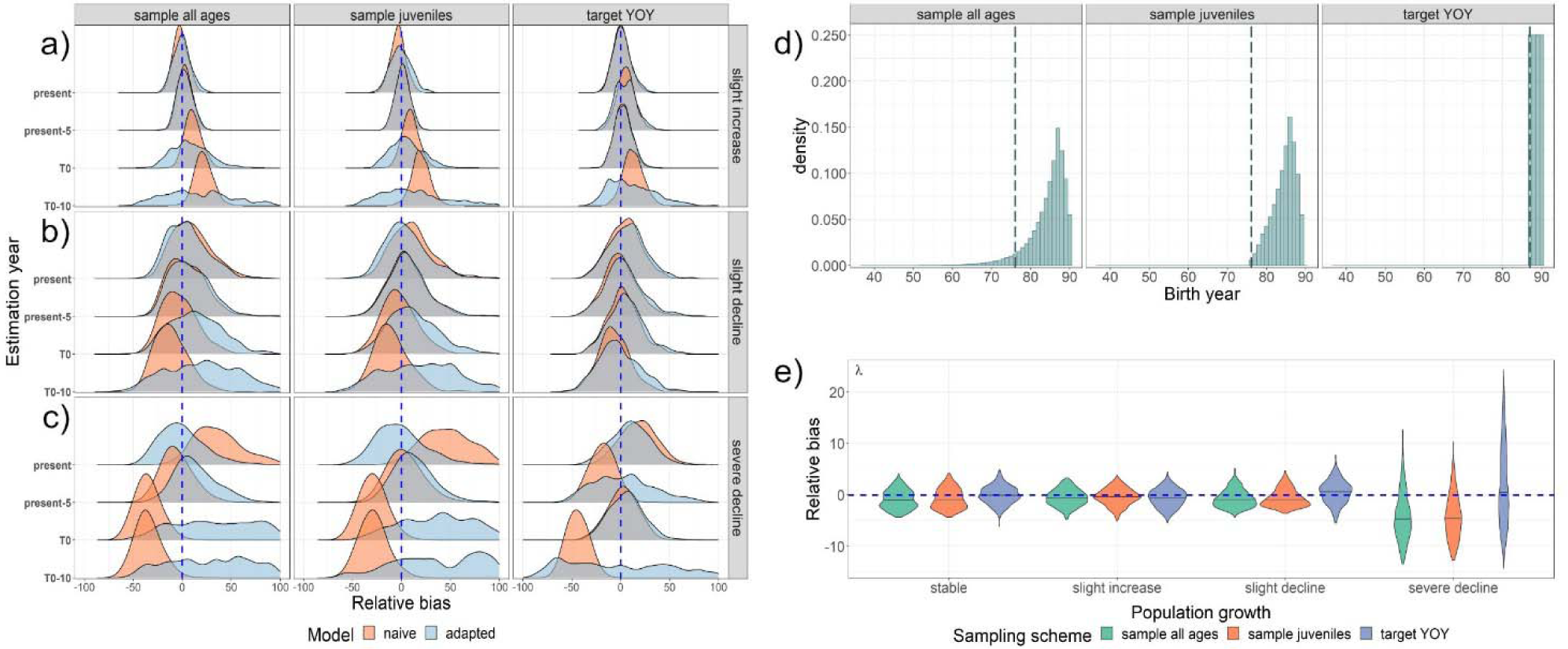
Performance of CKMR models when confronted with a changing population **a-c)** Relative bias of CKMR abundance estimates for mature females (*N*_♀*t*_) when applied to populations experiencing variable degrees of population growth or decline. Plots are split by sampling scheme (column), population growth pattern (row facet) and the year to which abundance estimates were targeted (year *t*). The dashed vertical blue line represents 0% relative bias. Scenarios assessed had population growth as **a)** slightly increasing (+1% per year), **b)** slightly declining (−1% per year), or **c)** severely declining (−5-10% per year over the final 10 years). Two different models were fit to 500 simulated populations for each scenario: a naïve model without a parameter for population growth (red) and an adapted model that included a parameter (λ) for population growth (blue). Plots were truncated at +/-100% for visualization because there were long tails of positive bias for the 10 year past scenarios. Note that for the target YOY sampling scenario, *t_0_* (the first instance of *y_j_* in the dataset) occurred three years in the past, making this the only scenario where an estimation year of *t_0_*occurred more recently than when the estimation year was “present-5”. **d)** Summary of age distribution of samples for all three sampling scenarios. The dashed vertical line represents *t_0_*, which varied depending on the ages sampled. Many more samples were drawn during year *t_0_*for the target YOY sampling scheme than for any other. **e)** Relative bias of λ estimates over the modeled time period.

Estimates of λ were highly correlated with abundance (Table 3; Appendix S1: S1.3) and were mostly unbiased for scenarios that involved a population that was monotonically increasing, declining, or stable (Figure 3e); however, the model tended to underestimate λ when the population began a severe decline during the modeled time period (“severe decline” scenario). Estimates of □ were unbiased regardless of population trend but varied with the number of age classes sampled (Appendix S1: Figure S3b) and were highly correlated with abundance in the scenario that only included four cohorts (target YOY; Table 3). Combined, these results suggest that a CKMR model that is adapted for population growth can give unbiased year-specific abundance estimates across a range of scenarios, while estimates of population trend should be interpreted with caution (see also section 3.5 and section 4.4 below for considerations to improve trend estimation).

**Table 3:** Mean cross-correlation values between female abundance at *t*_0_ (*N*_♀_*_(t0)_*) and survival (□) or population growth rate (λ) from population growth simulations (Table 1, scenario 2.1 – 2.4).

| Parameter | □ | $\lambda$ | $t_0$ | Sampling scheme |
| --- | --- | --- | --- | --- |
| $N_{\varphi(t_0)}$ | 0.77 | -0.13 | 88 | target.YOY |
| $N_{\varphi(t_0)}$ | 0.34 | -0.76 | 77 | sample all juveniles |
| $N_{\varphi(t_0)}$ | 0.28 | -0.80 | 77 | sample all ages |

### 3.3 Intermittent breeding

When a CKMR model that is fully naïve to intermittent breeding (Figure 4, orange) was applied to data from populations with females that bred on a consistent multiennial schedule, estimates of female (Figure 4a) and male (Figure 4b) abundance were positively biased. Males did not breed on a multiennial schedule, but they did share the survival parameter (□) with females, and this parameter was also overestimated with the naïve model (Figure 4c).

**Figure 4:**
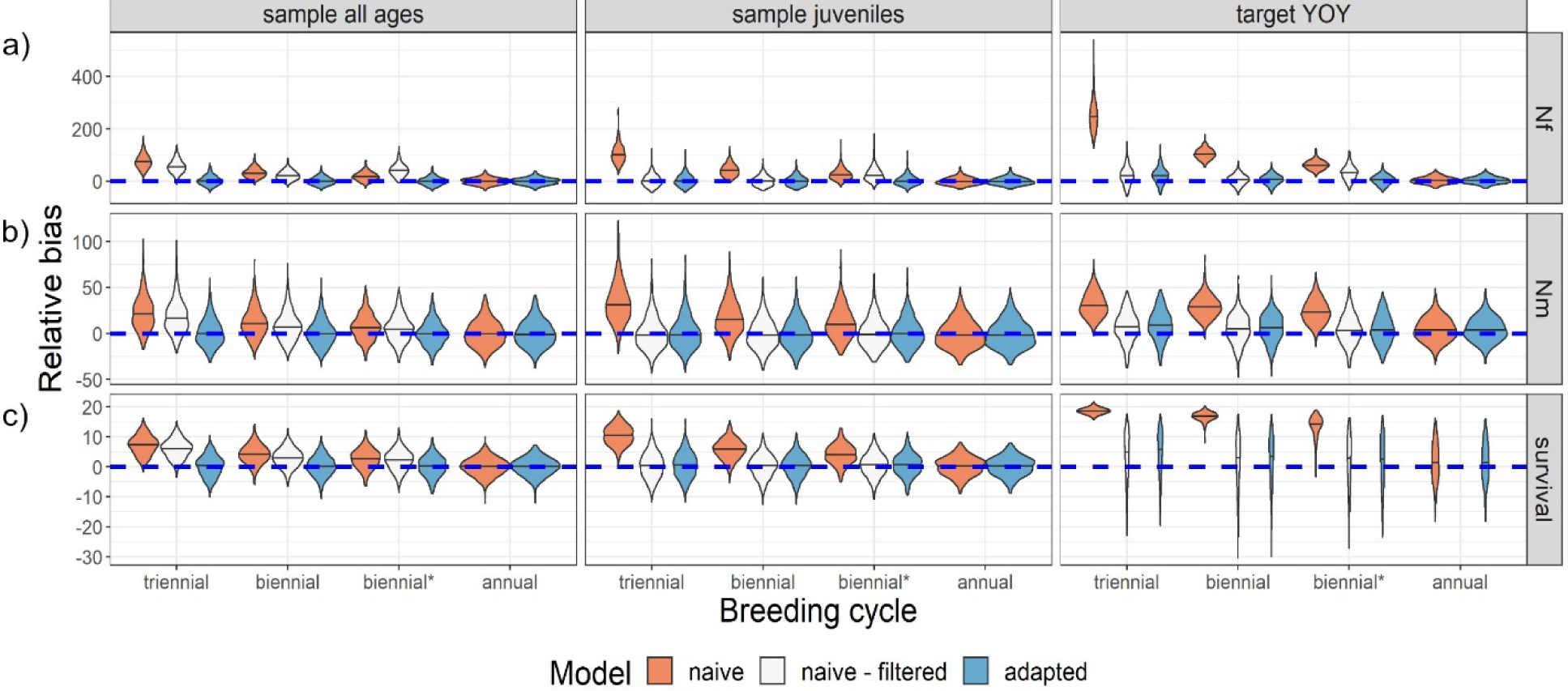
Relative bias of parameter estimates when naïve and adapted CKMR models were fit to samples from a population that breeds intermittently. The “naïve” model references a model that was included off-cycle comparisons, while the “naïve – filtered” model excluded off-cycle comparisons. The reference year (*t_0_*) was set to the earliest instance of *y_j_* and abundance was estimated in the present via Eq. 3 and compared to the true abundance in that year. For the “naïve – filtered” scenario, the quantity estimated by the model was the effective female breeders in year *t* (*Ñ_t_*), while the “naïve” and “adapted” models estimated total females (*N*_♀*t*_, or *Nf*). We derived total *Nf* for the “naïve – filtered” scenario by multiplying *Ñ_t_* by the breeding cycle (2 for biennial breeders and 3 for triennial breeders). On the x-axis, “biennial*” refers to a situation where 10% of biennial breeders bred off-cycle and 10% of on-cycle females failed to breed each year. **a)** Relative bias of abundance estimates for females (*Nf*). **b)** Relative bias of abundance estimates males (*Nm*). **c)** Relative bias of survival estimates.

When the pairwise comparison matrix was filtered to remove off-cycle comparisons before fitting a model that was otherwise naïve to intermittent breeding (Figure 4, white), estimates of abundance and survival were unbiased for models that only included HSPs (“sample juveniles” and “target YOY” scenarios), but only if 100% of females bred on the same schedule. When off-cycle breeding was introduced (biennial* scenario), estimates of female abundance were positively biased. When the model also included kinship probabilities for parent-offspring pairs (“sample all ages” scenario), estimates of female abundance, male abundance, and survival were all positively biased with the “naïve – filtered” model, reflecting the fact that the two kinship probabilities refer to different quantities (see section 4.2 and Appendix S1:S1.5).

Parameter estimates were generally unbiased with the model that was adapted for intermittent breeding (Equation 7), including the scenario with 100% annual breeders, in which case the naïve and adapted models performed identically (the “naïve – filtered” approach was not tested in this scenario because with annual breeding there were no off-cycle comparisons to remove). When we compared estimates of ψ to the realized proportion of HSPs that came from on-cycle females (the ‘true ψ’ in the simulated data), estimates of ψ were unbiased when all females bred on a multiennial cycle, with or without stochastic off-cycle breeding (Figure S5), though we note that when the multiennial model was applied to a population that bred annually, ψ had a non-unique solution (but estimates of other parameters were unbiased; see Appendix S1: S1.5 for further discussion about considerations surrounding intermittent breeding). Taken together, these results demonstrate that our model that accounts for multiennial breeding can accommodate variable or unknown breeding schedules and produce unbiased estimates of abundance and survival for populations that breed annually, biennially, or triennially, with or without instances of off-cycle breeding.

### 3.4 Aging uncertainty

When ages were misassigned to samples, older individuals were far more likely to be assigned to the wrong cohort (Figure 5a). The probability of age misassignment roughly corresponded to the slope of the von Bertalanffy growth curve, with the probability of age misassignment being greatest as the curve approached its asymptote. Consequently, when sampling was targeted to YOY, individuals were far more likely to be assigned to the correct cohort (Figure 5a, right).

**Figure 5:**
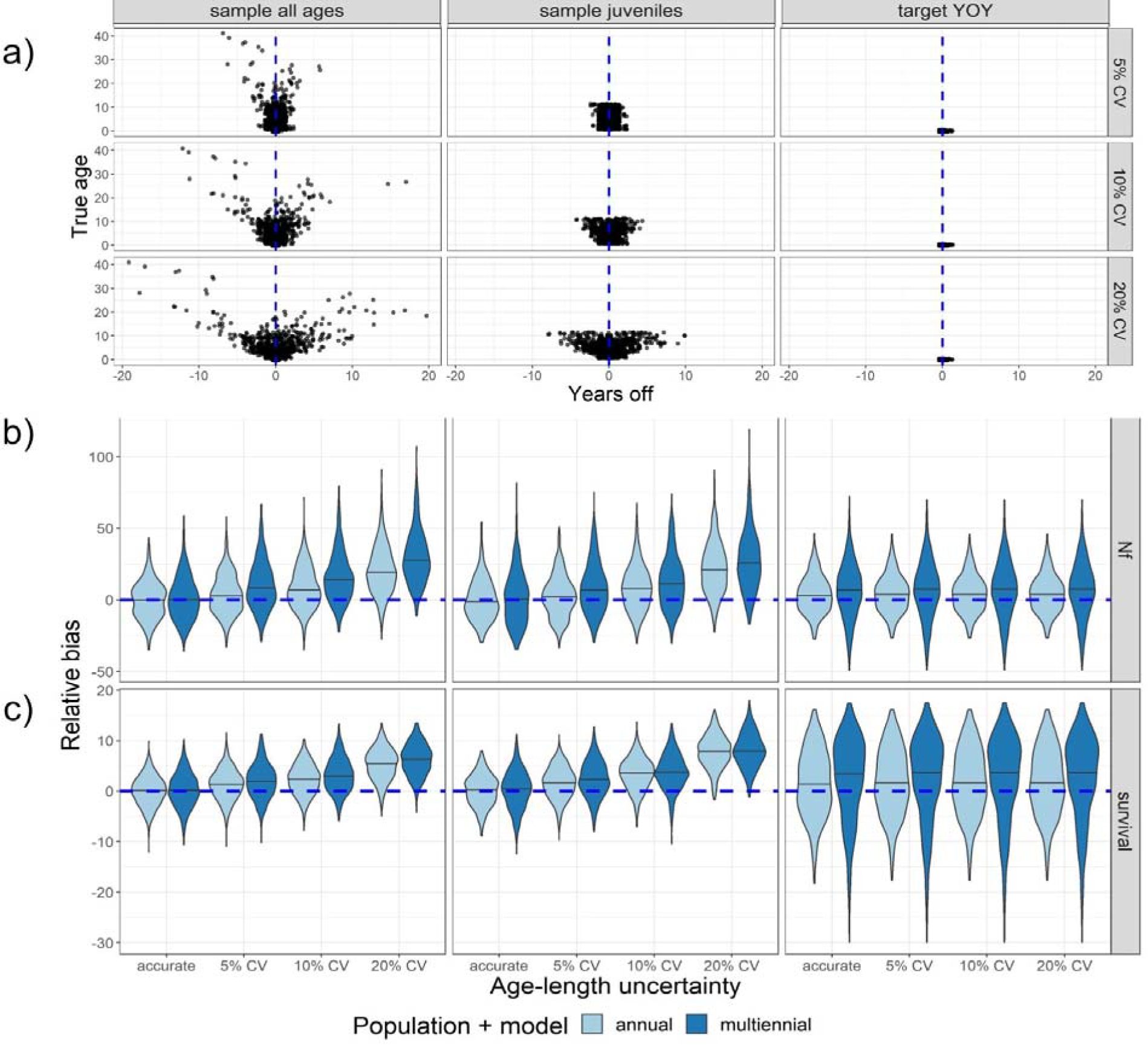
Effect of aging error on CKMR parameter estimates. **a)** Amount of error introduced per age for one of the 500 iterations that were run to test the effects of age misassignment on CKMR parameter estimates. The iteration represented here was chosen randomly and assumed to be generally representative of all 500 iterations. **b)** Relative bias of abundance estimates for females (*Nf*) when uncertainty was introduced to length-based age assignments. **c)** Relative bias of survival estimates when uncertainty was introduced to length-based age assignments. There was no intentional model misspecification in these simulations; rather, annual models were fit to populations that bred annually (light blue), while multiennial models were fit to populations that bred biennially (dark blue), thereby isolating the effects of aging error on the resulting bias.

When multiple age classes were represented in the data, bias accrued in estimates of female abundance (Figure 5b) and survival (Figure 5c) as the CV surrounding age assignment widened, regardless of whether we simulated a population that bred annually or biennially. Targeted sampling of YOY showed a different trend: the probability of misassigning an age-0 individual to the wrong cohort was very low, so increasing the CV on length-based age assignment did not affect the bias of parameter estimates. These results confirm that reliable aging is a key component of CKMR and that targeted sampling of age classes that can be reliably aged can improve estimation when accurate aging for other age classes is challenging.

### 3.5 Application to lemon sharks

#### 3.5.1 Bimini lemon shark simulations

When we simulated a small population that resembled the population of lemon sharks at Bimini, Bahamas, and then heavily sampled that population and used all samples for construction of the pairwise comparison matrix, results varied depending on which window of data was used (Figure 6 a-c). When mortality was reduced in the 90-year simulation and the population was growing (years 70-79), year-specific estimates of female abundance, abundance trend (λ), and survival (□) were mostly unbiased for the five-year window and the scenario that included all available samples (Figure 6a-c). As mortality increased and the population stabilized (years 80-84), bias began to accrue for the scenario that included all available samples. Bias continued to rise for this scenario when mortality was increased to produce a declining population (years 85-90). In contrast, results from the five-year window of samples were unbiased whenever the sample window spanned a period that included a consistent population trend: each time a shift in population trend occurred (years 80 and 85), the five-year window of samples produced biased parameter estimates for the following year, but that bias was reduced as the population trend stabilized. The three-year window of samples produced parameter estimates that were both biased and imprecise, suggesting that more than three cohorts are needed to produce reliable parameter estimates for a population that breeds biennially. All sample windows estimated abundance at an absolute scale within an order of magnitude of the real population size (see Appendix S1, Figure S6 for an illustrative example).

**Figure 6:**
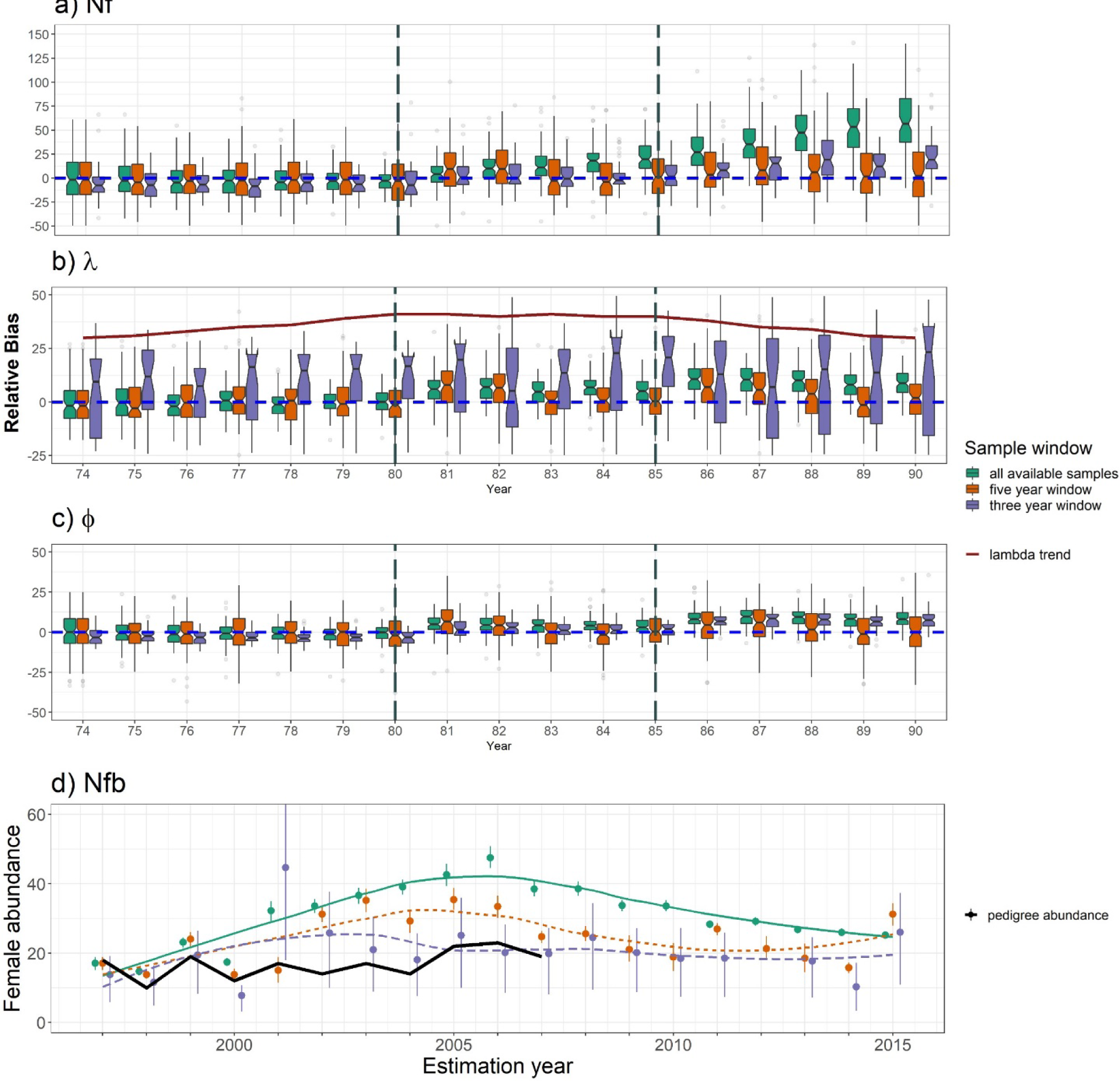
Time series of CKMR parameter estimates for simulated (**a-c**) and real (**d**) female lemon sharks at Bimini, Bahamas using all samples collected before the estimation year (green; solid trendline), all samples collected in a five year window prior to the estimation year (orange; dotted trendline) and all samples collected in a three year window prior to the estimation year (purple; dashed trendline). **a-c)** Relative bias from 100 distinct population simulations and model fits. **a)** Relative bias of abundance estimates for adult females (*Nf*) in each year of the time series. **b)** Relative bias of λ estimates relative to the observed population growth rate in the associated estimation year. **c)** Relative bias of survival (□) relative to the observed survival rate in the associated estimation year. **d)** Abundance estimates for breeding females (*Nfb*) in the North Bimini Lagoon using real genetic data, derived from estimates of *Nf* using Eq. 6. Points represent the median of the posterior distribution, and error bars reflect the 95% highest posterior density interval (HPDI). The trend is visualized using a loess regression. The black line labeled as “pedigree abundance” is a time-series of abundance estimates for *Nfb* that was independently derived for the population by Dibattista et. al. (2011).

When we retained just one full sibling from each mother/father pairing, parameter estimates showed similar tendencies as when the full dataset was used, with the exception that estimates of λ were less biased for the three-year window (see Appendix S1: S1.6; Figure S7 a-c). Combined, these results demonstrate that CKMR can reliably estimate abundance within an order of magnitude for a small population that is heavily sampled, while inference of trends – whether through a time-series of female abundance or estimation of λ – varies depending on the window of samples included.

#### 3.5.2 Bimini lemon shark data

Application of our multiennial CKMR model to real data from Bimini lemon sharks showed a parabolic abundance trend and very low abundance regardless of whether we retained full siblings (Figure 6d, Table 4) or filtered them (Appendix S1: Figure S7d, Table S3), and whether the datasets were downsampled (Appendix S1: Table S4, Table S5, Figure S8) or not. These trends were consistent regardless of which window of samples was used, though the timing of peak abundance varied (see Appendix S1: S1.6 for more discussion). Estimates of female breeders (*N*_♀*b*_) were close to, but slightly higher than, estimates independently obtained by <u>DiBattista et al. (2011)</u> using a pedigree-based approach.

**Table 4:** Year-specific abundance estimates for *N*_♀*b(t)*_ from the real dataset for Bimini lemon sharks when all individuals were kept from each mother/father pairing.

| Estimation year | All available samples |  |  |  | Five-year window |  |  |  | Three-year window |  |  |  |
| --- | --- | --- | --- | --- | --- | --- | --- | --- | --- | --- | --- | --- |
|  | Median | 95% HPDI | Total samples | MHSPs | Median | 95% HPDI | Total samples | MHSPs | Median | 95% HPDI | Total samples | MHSPs |
| 1997 | 17 | (15-19) | 285 | 807 | 17 | (15-19) | 285 | 807 | **14** | **(6-19)** | **257** | **570** |
| 1998 | 15 | (14-16) | 367 | 1322 | 14 | (13-15) | 354 | 1148 | 12 | (5-16) | 245 | 324 |
| 1999 | 23 | (22-24) | 489 | 2279 | 24 | (22-26) | 461 | 1848 | 19 | (8-27) | 312 | 501 |
| 2000 | 17 | (16-19) | 569 | 3204 | 14 | (12-16) | 447 | 1623 | 8 | (3-11) | 284 | 566 |
| 2001 | 32 | (29-35) | 651 | 3623 | 15 | (11-19) | 474 | 1347 | 45 | (18-65) | 284 | 175 |
| 2002 | 34 | (32-36) | 722 | 4380 | 31 | (29-34) | 437 | 1213 | 26 | (10-38) | 233 | 166 |
| 2003 | 37 | (34-39) | 796 | 5298 | 35 | (32-39) | 429 | 933 | 21 | (9-31) | 227 | 227 |
| 2004 | 39 | (37-41) | 886 | 6219 | 29 | (26-32) | 397 | 915 | 18 | (8-26) | 235 | 249 |
| 2005 | 43 | (39-46) | 1038 | 7489 | 35 | (32-39) | 469 | 1228 | **25** | **(10-36)** | **316** | **309** |
| 2006 | 47 | (45-51) | 1148 | 8825 | 33 | (30-37) | 497 | 1213 | **20** | **(8-28)** | **352** | **358** |
| 2007 | 38 | (36-41) | 1268 | 10745 | 25 | (23-26) | 546 | 1743 | **20** | **(8-27)** | **382** | **681** |
| 2008 | 39 | (36-41) | 1328 | 11616 | 26 | (23-28) | 532 | 1536 | 25 | (9-34) | 290 | 212 |
| 2009 | 34 | (32-35) | 1451 | 13461 | 21 | (18-25) | 565 | 1931 | 20 | (9-27) | 303 | 555 |
| 2010 | 34 | (32-35) | 1501 | 14339 | 19 | (15-23) | 463 | 1144 | 18 | (7-27) | 233 | 114 |
| 2011 | 28 | (27-29) | 1626 | 17072 | 27 | (25-29) | 478 | 1941 | 19 | (7-26) | 298 | 582 |
| 2012 | 29 | (28-30) | 1637 | 17144 | 21 | (18-25) | 369 | 813 | 168 | (26-442) | 186 | 15 |
| 2013 | 27 | (26-28) | 1716 | 19006 | 19 | (14-23) | 388 | 1281 | 18 | (7-25) | 215 | 410 |
| 2014 | 26 | (25-27) | 1749 | 19519 | 16 | (14-17) | 298 | 603 | 10 | (3-17) | 123 | 27 |
| 2015 | 25 | (24-26) | 1847 | 21700 | 31 | (28-34) | 346 | 1066 | 26 | (11-37) | 210 | 217 |

*Note:* MHSPs refers to the number of maternal half-sibling pairs identified. Bold reflects instances that included at least one parameter where $1.01 < \text{Rhat} < 1.02$ .

Estimates of adult survival ([£) were generally high for the full and downsampled datasets, but varied with the sample window (Appendix S1: Figure S9) and showed more variability when the dataset was both filtered for full siblings and downsampled (Appendix S1: Figure S8b). The three-year window of samples gave rise to survival estimates that were highly correlated with estimates of abundance (Table 5) and did not vary across years (Appendix S1, Figure S9), again suggesting that the three-year window is too short a time period for reliable estimation of survival rates for populations that breed biennially. Overall, regardless of how the data were subset, our results align with other studies that suggest low abundance (DiBattista et al. 2011) and high survival rates (White et al. 2014) of adult females at Bimini; however, we also note that estimation of abundance trend and survival were correlated with the number of cohorts included in the analysis and the mortality regimes that the population experienced.

**Table 5:** Cross-correlation among parameter estimates following application of CKMR to Bimini lemon sharks.

| Parameter | $N_{\varphi(t\theta)}$ | $\varphi$ | $\lambda$ | $\psi$ | Time window |
| --- | --- | --- | --- | --- | --- |
| $N_{\varphi(t\theta)}$ | 1.00 | 0.13 | -0.84 | -0.08 | all available samples |
| $\varphi$ | 0.13 | 1.00 | 0.32 | -0.10 | all available samples |
| $\lambda$ | -0.84 | 0.32 | 1.00 | 0.02 | all available samples |
| $\psi$ | -0.08 | -0.10 | 0.02 | 1.00 | all available samples |
| $N_{\varphi(t\theta)}$ | 1.00 | 0.29 | -0.73 | -0.18 | five-year window |
| $\varphi$ | 0.29 | 1.00 | 0.19 | -0.11 | five-year window |
| $\lambda$ | -0.73 | 0.19 | 1.00 | 0.09 | five-year window |
| $\psi$ | -0.18 | -0.11 | 0.09 | 1.00 | five-year window |
| $N_{\varphi(t\theta)}$ | 1.00 | 0.83 | -0.43 | -0.54 | three-year window |
| $\varphi$ | 0.83 | 1.00 | -0.01 | -0.46 | three-year window |
| $\lambda$ | -0.43 | -0.01 | 1.00 | 0.21 | three-year window |
| $\psi$ | -0.54 | -0.46 | 0.21 | 1.00 | three-year window |

*Note:* Reported values represent the average cross-correlation values over all years (1997-2015).

## 4. Discussion

Obtaining unbiased estimates of abundance is a central challenge for effective conservation and management of many threatened and exploited populations and is especially pertinent for populations of low density and highly mobile species where effective sampling of adults is impractical. Our simulation results broadly concur with recent work supporting CKMR as a promising approach to estimate abundance and survival in data-limited circumstances, but emphasize the critical need to adapt CKMR models adequately to accommodate population dynamics and life history traits that violate the assumptions of a simple base-case model. Further, although we confirm sensitivity of CKMR to aging error, we also find that bias in parameter estimates can be mitigated by sampling as few as four cohorts that can be reliably aged, providing options for applying the method when accurate aging is difficult or when long-term sampling is impractical. Our application to lemon sharks in Bimini, Bahamas demonstrates that CKMR is a flexible framework that can be used to estimate abundance and survival of breeding adults when only juveniles are available for sampling. Taken together, the results of our application of CKMR to simulated and real populations with different population sizes, trends, and breeding schedules support the recognition of CKMR’s immense potential for monitoring populations of low density and highly mobile species, while also highlighting several promising avenues for future research.

### 4.1 Accounting for population growth/decline

A simple base-case CKMR model (e.g., Eqs. 1 and 2) estimates adult abundance over the modeled period by assuming that population dynamics are stable and consistent over time. In cases with sex-specific or transient population dynamics, or if estimates of underlying population parameters are desired, population dynamics can be modeled with CKMR using latent variables. Alternatively, year-specific abundance estimates can be obtained by modeling each year independently, but this approach requires a particularly rich dataset (e.g., heavily sampled salmonids; Ruzzante et al. 2019, Marcy-Quay et al. 2020). As such, most data-limited situations will likely benefit from leveraging all available data for a single abundance estimate. In such cases, specifying an exponential growth model where population dynamics are broadly captured in a parameter like λ allows data to be shared across cohorts to produce a single estimate of abundance for a specified reference year *(t_0_*). Then, abundance in any modeled year can be derived from estimates of *N_(t0)_* and λ. In practical applications of CKMR, knowledge of a species’ life history in combination with Leslie matrix simulations can help inform a prior on λ to improve precision of parameter estimates. If a fishery-independent index or fishery-dependent CPUE index (from a fishery whose operations have been relatively constant) exists over the modeled period, then trend data could also be integrated into the model via specification of the prior on λ.

Including too many age classes in the data may hinder inference of abundance trends if a population’s trajectory shifts during the modeled time period, as estimates of λ will represent an average of those trajectories. One way to mitigate this averaging effect is to subset the dataset for smaller time windows over which averaging λ has a less pronounced effect. This approach sacrifices precision and may not be possible in many data limited situations; even in a data-rich scenario, the time window must be thoughtfully calibrated, as using too small a window provides limited data to the exponent on □, resulting in a high degree of correlation between □ and *N* (e.g., the target YOY scenario in Table 3 and the three-year window in Table 5). On the other hand, our results suggest that using too large a window will result in an abundance trend that lags behind the real trend if the real trend shifts during the modeled period. For our Bimini lemon shark simulations, a five-year window produced a reasonable balance of precision and accuracy for parameter estimates and also for reconstructing an abundance trend for a population that experienced multiple mortality regimes during the modeled period. In application to real data sets, one would not know the accuracy associated with a given window-size, but the pattern of lag in estimated population trends as the window length is expanded may give some indication of where shifts in the trajectory have occurred. Alternatively, one could test for a quadratic trend by adding another parameter in the model for λ and then use model selection to determine the model that best captures the true population trend. Expanding CKMR to estimate population trend in a reliable and robust way is a ripe area for future research.

### 4.2 Intermittent breeding dynamics

When intermittent breeding coincides with a population that is most easily sampled during the juvenile life stage (e.g., when adults are not directly observed), our results indicate that abundance estimates derived from a naïve half-sibling CKMR model will be biased if all pairwise comparisons are included in the model. In contrast, in circumstances where all individuals breed on the same schedule such that there is no possibility of off-cycle breeding, then filtering the pairwise comparison matrix to remove off-cycle comparisons and fitting a half-sibling model that is otherwise naïve to intermittent breeding (e.g., Eq. 4) can give unbiased parameter estimates (Figure 4, white). Importantly, when off-cycle comparisons are removed, the non-breeding adults essentially become invisible to half-sibling CKMR (similar to infertile adults; see section 3.2 of Bravington et al. 2016a). As such, a naïve HS model that is filtered to remove off-cycle comparisons gives estimates of the number of effective female breeders (*Ñ*) in year *t* (Appendix S1: S1.4), a number that may be substantially different than the total number of adult females (*N*) in populations that breed intermittently. Without modification to the kinship probabilities, this difference precludes the use of POPs in the same model because PO CKMR gives estimates of *N* (note the bias for the “sample all ages” scenario in Figure 4). In addition, our simulations suggest that if off-cycle breeding produces HSPs that are separated by a birth year gap that does not align with the expected breeding cycle (e.g., when the birth year gap is odd for a population that breeds on a biennial schedule), then parameter estimates will be biased with a naïve model, whether off-cycle comparisons are retained in the pairwise comparison matrix or not. Overall, applying a model that is naïve to intermittent breeding to a population that breeds on a bi- or triennial schedule can produce unbiased parameter estimates if off-cycle comparisons are removed, but only in limited situations (e.g., when there is no off-cycle breeding).

The multiennial CKMR model presented here accommodates intermittent breeding via inclusion of the parameters Ψ and *a*, which assigns a non-zero probability to off-cycle comparisons without assuming the probability is the same as on-cycle comparisons, resulting in estimates of *N* rather than *Ñ* (see Appendix S1, S1.4 and S1.5 for further discussion). While the parameter Ψ can be estimated, *a* must be fixed to the expected breeding cycle. If the breeding cycle for a population is unknown, and if adults are not available for sampling, then it may be possible to estimate *a* from the distribution of birth year gaps among identified half-siblings (Waples and Feutry 2021). As a cursory example, if most HSPs were born in year gaps that are divisible by 2, then fixing *a* to 2 would be logical. However, in real populations, reproductive periodicity may be challenging to infer from the distribution of kin pairs, as environmental conditions may cause individuals to fail to breed one year and then breed off-cycle the next (Rivalan et al. 2005, Cubaynes et al. 2011, Morbey and Shuter 2013, Öst et al. 2018, Skjæraasen et al. 2020). In our lemon shark dataset, for example, 30% of adult mothers bred off-cycle at least once before returning to a biennial cycle. With elasmobranchs and other species that are difficult to age, aging error will further obscure inference of breeding schedule based on offspring birth years. Stochastic off-cycle breeding was not a problem for our multiennial model as long as there were no systemic differences in lifetime fecundity (Bravington et al. 2016a; see Appendix S1:S1.6 and Figure S10). Future work that adapts CKMR to estimate Ψ and *a* across a range of scenarios, including populations with mixed mating schedules (Walker 2007, Driggers and Hoffmayer 2009, Higgs et al. 2020), would further expand the potential of CKMR to illuminate aspects of population breeding dynamics.

#### 4.3 Aging error

CKMR depends heavily on accurate cohort assignment, which can be very challenging for many species, including elasmobranchs. Our results confirm that age misassignment can substantially bias CKMR parameter estimates. A hierarchical model that accounts for aging error may help alleviate this issue, but such a model would require some estimate of the probability of age misassignment (Hirst et al. 2004, Schwarz and Runge 2009) and selectivity (Henríquez et al. 2016, Francis 2016), and such data may not be available in data-limited situations. Estimating the probability of age misassignment is not a trivial task, even for species with well-established aging methods (e.g., teleosts, <u>O’Sullivan 2007)</u>. For species that are challenging to age, substantial upfront effort may be required to calibrate aging methods and estimate the degree of error present. For example, patterns of DNA methylation can be used to estimate age (Jarman et al. 2015) and these data can be obtained from the same tissue samples used for kinship assignment in CKMR. However, epigenetic clocks are taxa-specific, and the discovery of informative biomarkers requires calibration using representative samples of known ages, which may be arduous to obtain in their own right (Polanowski et al. 2014, Beal et al. 2022). It is wise, therefore, to consider how samples will be aged and how much error there is likely to be prior to embarking on a large-scale CKMR study.

In cases where only YOY can be reliably aged, our results show that CKMR can generate reliable abundance estimates from targeted sampling of as few as four cohorts of YOY, even for a population that breeds bi- or triennially, though estimates of survival will improve as more cohorts are added. If mature individuals are also available to sample – e.g., when visiting a nursery site to breed – then sampling potential parents as well as YOY can enable the use of POPs in the likelihood and improve precision of parameter estimates. Aging error in this case would be less critical for adults as long as maturity can be confirmed in the year of sampling, though care must be taken to ensure that potential parents and offspring are sampled independently, as parameter estimates will be biased if the probability of sampling a parent is correlated with the probability of sampling its offspring (Bravington et al. 2016a).

### 4.4 Population dynamics and abundance of lemon sharks in Bimini

Our application of CKMR to Bimini lemon sharks highlights the flexibility and potential of CKMR for long-term monitoring of populations of low density highly mobile species with geographically distinct life histories. Estimates of abundance from CKMR suggest that a very small number of female lemon sharks give birth at the North Bimini Lagoon during each biennial breeding cycle (Figure 6d, Table 4). These results align with a previous study that reconstructed a pedigree for the population and identified the number of adults that successfully bred at the North Island each year between 1995-2007 (DiBattista et al. 2011). In both cases, the number of females that gave birth at the North Island during this time period was estimated to be very small (<50 per year), with an increasing abundance trend through ∼2006. At some point after or around 2006, results from CKMR suggest that the number of females using Bimini for breeding began to decline. Intense dredging and mangrove deforestation took place around the North Bimini Island in March 2001 in preparation for development of a mega-resort (Jennings et al. 2008). Although the number of breeding females at the North Island counterintuitively increased immediately after the disturbance (DiBattista et al. 2011), there was a transient drop in the survival rates of age 0 and age 1 individuals, though the degree to which juvenile mortality was affected is debated (Jennings et al. 2008, DiBattista et al. 2011). These cohorts would have reached maturity and begun returning to Bimini for reproduction around 2011, which may explain the decreasing trend around that time (Figure 6d). All sampling windows and methods of downsampling showed a parabolic abundance trend over the time series, though the stationary point (where population size was stable) varied depending on the window.

Although our results closely resemble those reported in DiBattista et al. (2011), we note that our abundance estimates from CKMR were slightly higher. The degree to which our results differed depended on whether we included full siblings in the analysis and whether we used the full dataset or a downsampled dataset (see Appendix S1: S1.6 for more discussion on CKMR with small populations). Abundance estimates were generally similar (<50 breeding females) across the datasets we tested, except for a few instances when sampling was constrained to a three-year window. When a population breeds biennially, sampling three years only includes one year gap with possible positive comparisons (years one and three), which provides very limited information to the exponent on □ and impedes its estimation (see Appendix S1: Figure S9), as well as the estimation of other parameters (note the high correlation with abundance in Table 5).

Though all three windows of samples we tested (three-year, five-year, all available) suggest the population of breeding females at Bimini is small, the five-year window produced the least biased parameter estimates in simulation, and application to the real data resulted in estimates that aligned more closely with the estimates of Dibattista et al. (2011) than when all samples were used. More complex models (e.g., that allow for quadratic abundance trends) would likely improve performance of models that leverage long time-series of data; in the absence of such a model, calibrating the time window of samples to correspond to likely abundance trends can help alleviate the averaging effect of assuming exponential growth/decline over long time periods.

Adapting CKMR to produce more reliable parameter estimates for small populations will require additional work (Bravington et al. 2016a); however, when dealing with abundance estimates that are small enough to cause such issues, the practical implications of this known bias are likely minimal. Discarding estimates from the three-year window of samples, our model that retained full siblings in the dataset estimated a maximum of 47 females visited the North Island across all 18 years of abundance estimation (Table 4, Figure 6d). Removing full siblings and downsampling the dataset produced slightly higher values but still did not exceed 70 females (Appendix S1: Figure S7, Table S3). These quantities are small enough that any additional mortality would likely threaten the sustainability of this portion of the population.

### 4.5 Implications for sampling design

We have shown that application of HS CKMR to long-lived species can generate reliable estimates of abundance – on its own, or in conjunction with PO CKMR - from a limited number of cohorts when aging is reliable; however, estimates of survival (⍰) were more reliable when more cohorts were included in the dataset across our simulations. A dataset that spans enough cohorts to reliably estimate parameters beyond abundance can be obtained by intensely sampling multiple age classes over a small number of years, or by long-term sampling of nursery areas. The former would require reliable aging of all sampled age classes to avoid biased parameter estimates, especially for models that incorporate half-sibling kinship probabilities where estimates of survival and abundance both depend on the birth year of sampled individuals. The latter - long-term sampling of nursery areas - represents a promising method for monitoring low density highly mobile populations, especially in circumstances where aging error is likely for older age classes.

We are not the first to suggest that CKMR benefits from focusing sampling efforts on individuals that can be reliably aged (Trenkel et al. 2022). Our results expand on this idea by demonstrating that CKMR can produce robust abundance estimates from as few as four cohorts, though estimates of survival will be less reliable as fewer age classes are included. In cases where sampling of juveniles is focused on nursery areas, sufficient biological knowledge to determine the scope of inference for CKMR will be required; if the target population uses multiple nursery areas, then sampling multiple nurseries can allow the model to estimate demographic connectivity (Patterson et al. 2022a). If sex-specific population dynamics are present, as with Bimini lemon sharks, the associated CKMR model should account for this and estimate parameters separately for each sex or focus solely on the sex for which the scope of inference is well-understood, as we did with Bimini lemon sharks.

One of the more exciting aspects of CKMR is its potential to generate rapid estimates of adult abundance without sampling a single adult (see Patterson et al. 2022a for an applied example). Our results confirm that a sampling program that can procure as few as four or five reliably aged cohorts can be used in combination with half-sibling CKMR to produce robust estimates of present-day abundance as well as reasonable estimates of survival. In circumstances where a genotyping panel, workflow for assigning kinship, and appropriate CKMR model are already developed for a population, contemporary abundance estimates could conceivably be obtained within weeks of sampling. As such, CKMR can offer a rapid and cost-effective method for population monitoring in real time following an initial investment in the laboratory and analytical workflows.

## Conclusion and Future Directions

CKMR is a powerful tool for estimating population abundance of species that have been historically difficult to assess. Reliable application of the method requires careful consideration of the relevant population dynamics matched to an appropriate sampling scheme. Here, we have identified a set of factors that must be considered for robust application of CKMR, proposed methods for accounting for them, and highlighted areas in need of further research. Specifically, we found that a half-sibling focused CKMR model can produce robust abundance estimates from as few as four or five cohorts, while reliable estimates of survival will likely require more data. Monotonic abundance trends can be dependably inferred by incorporating a simple exponential growth model; however, more complex trends will require further model development or, at a minimum, deployment of a sliding window of samples, which prevents long-term averaging of λ and obfuscation of transient dynamics.

When ages are prone to misassignment, focusing sampling efforts on individuals with known ages (e.g., YOY), or subsampling for these individuals if the dataset is sufficiently rich, can alleviate bias in parameter estimates, particularly abundance. Long-term monitoring of highly mobile species can be enhanced by CKMR via sampling of nursery areas when one or both sexes are philopatric, and can provide estimates of present-day abundance and abundance trends for adults that visit the nursery area without directly sampling a single adult. Overall, this study highlights the sensitivity of simple base-case CKMR models to assumptions about population dynamics and sampling, while also demonstrating that the CKMR framework is easily adaptable to accommodate these factors, making it a promising tool for integration into long-term monitoring programs.

## Acknowledgements

The authors thank Paul Conn and Shane Baylis for early guidance with regards to CKMR model construction and individual-based simulation during the initial phase of the project, and Andy Lin and Patrick Sadil for assistance with code that was integrated into construction of the pairwise comparison matrix. We are grateful to Chris Sutherland for invaluable support during model validation, and to Mark Bravington, Mark Maunder, Kevin Piner, and Steve Teo for conversations and suggestions that guided our sensitivity tests. The authors are exceedingly grateful to Paul Conn and an anonymous reviewer for insightful comments on a prior version of this manuscript, and members of the Molecular Ecology and Conservation Lab at UMass Amherst provided helpful feedback on earlier drafts of this manuscript. Thanks to the hundreds of volunteers and staff at the Bimini Biological Field Station that sampled and tagged lemon sharks. This work was completed using resources provided by the University of Massachusetts’ Green High Performance Computing Cluster (GHPCC). Funding for this project was provided to JDS by the American Fisheries Society via the Steven Berkeley Conservation Fellowship, the Save Our Seas Foundation, and the UMass Amherst Organismic and Evolutionary Biology program.

## Appendix S1

### S1.1. Constructing the pairwise comparison matrix

Close-kin mark-recapture models use a “pseudo-likelihood” framework to estimate population parameters. Whereas a full likelihood would derive probabilities of kinship from the population’s full genealogy (which is infeasible in most cases), a pseudo-likelihood works by specifying a probability of kinship for each pair of sampled individuals. Assuming each pairwise comparison is independent, the pseudo-likelihood approximates the full likelihood and can be used in its stead. This equivalence is central to CKMR and requires thoughtful construction of the full pairwise comparison matrix that is used as the input to the CKMR model.

The full pairwise comparison matrix contains kinship probabilities for each pair of sampled individuals. This matrix – which is likely to be tens of thousands or millions of rows depending on sample size – can be condensed into fewer rows by grouping comparisons based on the relevant covariates for the CKMR model (e.g. by reference year, birth year gap, and reference year gap; Table S2). The relevant covariates vary based on the parameterization of the CKMR model, which means that each model will allow for different degrees of grouping; the more the data can be grouped, the faster the model will run at the expense of accounting for individual variation.

Besides grouping the data – which will depend on how the model is constructed – there are various considerations to keep in mind when constructing the final pairwise comparison matrix for CKMR. First, comparisons with a zero percent chance of unveiling kin should be excluded. Alternatively, careful construction of the model to ensure that such comparisons are assigned zero probability of kinship in the model will suffice (but including both filters is optimal). For parent-offspring relationships, this means removing comparisons between offspring and other individuals that could not have birthed/sired them (e.g., because they themselves were not sexually mature or alive during the offspring’s birth year). For half-siblings, comparisons should be excluded if the individuals were born too far apart for a single individual to have birthed both of them. For instance, if a species is expected to live to a maximum age of 10 years old and is reproductively mature at age 3 (assuming knife-edge maturity for this example), then comparisons between individuals born more than 7 years apart should be excluded from the final pairwise comparison matrix. That said, in most applications of CKMR, the boundaries of life expectancy and reproductive maturity for a population will not be so clear-cut; as such, a probabilistic approach that is directly integrated into the CKMR model may serve better when the year gap is near the expected maximum.

Besides including impossible comparisons, an additional consideration for half-sibling CKMR is whether or not to include within-cohort comparisons. The rationale and consequences of this decision have been outlined elsewhere (Bravington et al. 2016a, Waples and Feutry 2021); for a CKMR model that incorporates half-sibling relationships and does not explicitly account for same-year comparisons, the most straightforward approach to this issue is to only retain cross-cohort comparisons in the final pairwise comparison dataframe.

Finally, it is worth mentioning that it can be easy to accidentally double-count comparisons; as such, the pairwise comparison matrix should be double-checked to ensure that each comparison is only included once. We have provided annotated R code that can serve as a starting point (see Data Availability statement).

### S1.2. Instances of aunt/niece pairs

Including aunt/niece (uncle/nephew, etc.) pairs as HSPs had a minimal effect on our results (Appendix S1, Figure S1a), and the small amount of bias that was introduced to the “sample all ages” scenario was rectified by including a filter that removed all half-sibling comparisons that spanned a year gap greater than the age of maturity (11 years). Aunt/niece pairs were rare in our simulated dataset (Appendix S1, Figure S1b), as were full siblings (Appendix S1, Figure S2). Conflating aunt/niece pairs with HSPs is likely to be a greater issue for circumstances in which full siblings are more prevalent and/or when the first age at reproduction is early enough that constraining year gap comparisons to the age at reproduction is impractical. For long-lived promiscuous species like lemon sharks, a simple year gap filter set to the first age of reproduction should suffice to remove most contaminating instances of aunt/niece pairs.

### S1.3. Accounting for a changing population

To interpret CKMR abundance estimates and appreciate its limitations, it is important to understand that *each* pairwise comparison includes a specific year (*y_j_*) for which abundance is estimated. Crucially, *y_j_* (the birth year of the younger individual) is different for each pairwise comparison; however, pairwise comparisons can be grouped based on relevant covariates, including *y_j_*. If there are enough comparisons that share the same *y_j_*, then it may be possible to produce multiple independent estimates of *N_(yj)_*, but doing so requires a very rich dataset. Instead, most cases of CKMR application will benefit from leveraging all available data for a single abundance estimate. This can be accomplished by specifying a population growth model that links each instance of *y_j_*. For example, we applied a simple exponential growth model to Scenarios 2, 3, and 4, as well as to the small population simulations and Bimini dataset. To link the exponential growth model to our CKMR model, we first selected a reference year (*t_0_*), which was the earliest instance of *y_j_* in each dataset (i.e., the second oldest individual used in the half-sibling probabilities). Then, for each pairwise comparison, abundance was estimated in *y_j_* and linked to *t_0_* via λ (see Equations 4 – 9). By including the parameter λ, all of the pairwise comparisons were leveraged to produce a single estimate for *N_(t0)_*. Then, *N_t_* was derived from estimates of *N_(t0)_* and λ using Equation 3.

### S1.4. Deriving Equation 7

Let *a* be the breeding interval, *ψ* the fraction of females that breed every *a* years, *N*_♀*t*_ equal the total number of mature females in the population in year *t*, and *Ñ_t_* equal the number of effective female breeders in year *t*. If we assume that *1/a* of the non-annual breeders produce pups each year, then

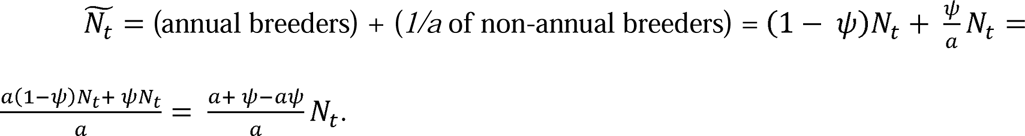

For individuals *i* and *j* born in years *y_i_*and *y_j_*, respectively, the probability that they have the same mother *d* is the product of the probability that *d* is mother of *i* and the probability that *d* is the mother of *j*. These probabilities are a function of relative reproductive effort and (when *y_j_>y_i_*) survival. We define relative reproductive effort as the reproductive output of mother *d* in a given year relative to the total reproductive output in that year. Assuming adult survival (*ϕ*) and fecundity (*f*) are constant across mature ages, then the probability that *i* and *j* are half-siblings is (from Bravington et al. 2016a):

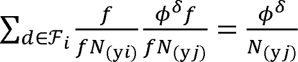

where the summation is across all breeding females at time *i*.

When breeding is intermittent, the denominators in the above equation (representing total reproductive output in a given year *t*) are the product of fecundity (*f*) and effective female breeders (*Ñ_t_*), and the summation is across the number of effective breeders at time *y_i_*. The first quotient inside the summation becomes 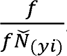. Given that a female was alive and reproduced in year *y_i_*, the probability that same female survived and is the mother of *j* is now dependent on the breeding cycle *a* and the fraction of the population that breeds every *a* years (*ψ*) or annually (1-*ψ*). There are two cases to consider, namely whether or not δ is divisible by *a*. When δ is divisible by *a*, then both annual and non-annual females breed; if δ is not divisible by *a* then only the annual breeders are capable of being the mother. Thus, we have the probability of a maternal half-sibling pair as

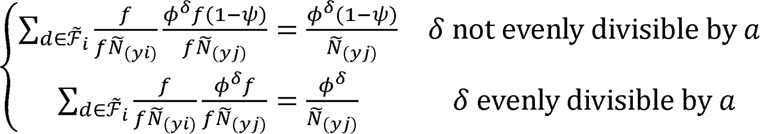

In the above, when δ is not divisible by *a*, the probability that mother *d* is an annual breeder is (1-*ψ*). When *δ* is divisible by *a*, the probability that mother *d* is either an annual or non-annual breeder is 1.

Replacing *Ñ_t_* with *N*_♀*t*_ we have equation (7):

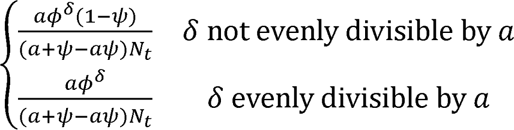

When *ψ*=1 and *a>1*, this yields the obvious result that 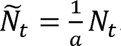. For example, if all females breed on a biennial cycle (i.e., *ψ* = 1 & a =2), then all gaps δ are multiples of 2 and we have:

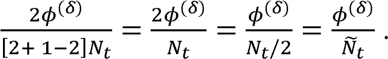

An alternative assumption for non-annual breeders is that *all* of them breed exactly every *a* years (rather than *1/a* of them breeding each year, as above). In this case, the number of effective breeders *Ñ_t_* in a given year depends on whether δ is divisible by *a*:

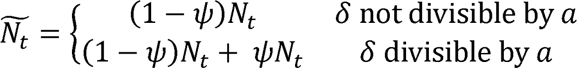

Thus, every *a* years, all mature females breed, and only (1 - *ψ*)mature females breed in the intervening years.

### S1.5. Intermittent breeding and systemic differences in lifetime reproductive output

Our simulation results show that multiennial breeding causes bias in juvenile-focused CKMR models that do not explicitly account for it. In circumstances where 100% of females breed on a multiennial cycle, the positive bias likely arises from the inclusion of pairwise comparisons that span year gaps without any HSPs (e.g., odd year gaps for biennial populations). Such comparisons can be filtered before model fitting to produce unbiased parameter estimates, including abundance, but the quantity (*N*_♀*b(t)*_) will be different than if off-cycle comparisons are included (*N*_♀*(t)*_). If off-cycle comparisons are excluded (or, alternatively, given a probability of 0 in the model), *N*_♀*(t)*_ can be derived by multiplying estimates of *N*_♀*b(t)*_ by the reproductive periodicity (e.g., 2 for biennial breeders). In most practical applications of CKMR, instances of off-cycle breeding and/or skipped breeding cycles are likely to result in some positive comparisons that do not align with the expected breeding cycle. Filtering comparisons from off-cycle breeding years would then result in loss of valuable data at best, and biased parameter estimates at worst (see Figure 4). In contrast, our multiennial model (Eq. 7) can leverage pairwise comparisons regardless of whether the birth year gap is on- or off-cycle and produce reliable estimates of total abundance and other parameters across a range of breeding cycles. Using this approach, *N*_♀*b(t)*_ can be derived from estimates of *N*_♀*(t)*_ using Equation 6, as we have done with Bimini lemon sharks.

Another circumstance that is related to differences in realized reproductive output and known to cause issues for CKMR is persistent individual differences in fecundity that give rise to lifetime differences in reproductive output. We confirmed this issue by fitting an annual and multiennial model to simulated populations with variable proportions of biennial and annual breeders *without* adjusting fecundity (Figure S10). This resulted in a population where annual breeders produced more offspring on average over their lifetimes, but this difference in reproductive output was not reflected in the kinship probabilities (Eq 7 assumes equal reproductive output among all females). Abundance estimates (primarily for females) using the multiennial model were more biased as a consequence of this unmodeled heterogeneity relative to when lifetime fecundity was constant across the population (Figure 4), though we note that the absolute bias was still less than 20% for two of the three sampling scenarios. Thus, our results confirm that unmodeled persistent differences in lifetime fecundity across different sectors of the population will, at a minimum, result in less reliable model performance, and we recommend future CKMR model development to accommodate persistent individual differences in fecundity.

### S1.6 CKMR with a small population

The pseudo-likelihood that CKMR employs has the properties of a full likelihood when sampling is sparse because each pairwise comparison among individuals is approximately independent (Bravington et al. 2016a). However, when CKMR is applied to very small populations (<∼100 individuals) and sampling is non-sparse the assumption of independence among samples is difficult to fulfill as capture of littermates, which may have similar genetic fitness and/or early-life history environments, becomes increasingly common. In such cases, it is important to account for or remove littermates to maintain independence among samples, since comparisons between littermates and a third animal are not independent (<u>Bravington et al.</u> <u>2016a)</u>.

In our application to simulated and real lemon sharks with very small population sizes (<100 total adult females), we found that applying CKMR to this small population without substantial modification to the likelihood produced estimates of abundance that were still very close to the true value. When we downsampled the real genetic dataset to 30% of the original sample size, the five-year window and the window that included all available samples produced parameter estimates that were similar to the full dataset but were predictably less precise (Figure S8, Table S4, Table S5). The combination of downsampling and limiting comparisons to a three-year window produced several instances of very few MHSPs arising from exceptionally small sample sizes, and the associated abundance estimates were unrealistic (Table S4, Table S5). Apart from those instances, the overall results remained similar whether we kept or excluded littermates, but retaining littermates produced parameter estimates that were less biased. We suspect that when populations are very small and heavily sampled such that pairwise comparisons are not independent, excluding littermates may skew the fecundity evident in the remaining samples, thereby increasing bias relative to when littermates are retained. By contrast, in larger populations retaining littermates would violate the assumption of non-independence for only a small subset of the potential comparisons, but a relatively large proportion of the identified true kin pairs. This could result in a large effect on parameter estimation, which may depend on relatively few kin pairs if sampling is sparse. Adapting CKMR for robust application to small populations is an active area of research that will likely require modifications to the statistical framework (<u>Bravington et al. 2016a)</u>, including adding a parameter for capture probability to PO kinship probabilities (see Section 2.4.1 of this paper). However, our results show that even in the absence of such modifications, CKMR can provide useful parameter estimates for small populations.

---

**Table S1.** Parameters for the data generating model (DGM). Values in bold produced, on average, a stable population size at equilibrium over the 90-year time course.

| Parameter | True value (DGM) |
| --- | --- |
| Female fecundity (total pups per year) | **5.8** (range: 2-9) |
| Breeding cycle | 1, 2, or 3 years |
| Initial adult population size | **1000** |
| Young-of-year survival (age 0) | **0.7** |
| Juvenile survival (ages 1-11) | **0.80** (range: 0.72 – 0.81) |
| Adult survival (age 12-50) | **0.825** (range: 0.75 - 0.83) |
| Age at maturity | **12** |
| Sex ratio | **50/50** |
| Maximum age | **50** |

**Table S2.** An example pairwise comparison matrix (subsetted for first 20 rows) used as input to a multiennial CKMR model (Eq. 6). <u>ref.year</u> is the birth year of the younger individual in the pairwise comparison (i.e., *y_j_*); <u>all</u> is the total number of comparisons that contain the same values for ref.year, mort.yrs, and ref.year.gap; <u>yes</u> is the total number of positive comparisons with those same values; <u>type</u> is either HS (half-sibling) or parent-offspring (PO); <u>parent</u> is mother or father; <u>ref.year.gap</u> is the reference year gap i.e. (*y_j_*– *t_0_*); <u>BI</u> refers to the breeding interval, which is either on-cycle or off (depending on *a*) for HS comparisons and irrelevant for PO comparisons. There are myriad ways to construct a pairwise comparison dataframe for CKMR; we found that producing the table below allowed us to store all comparisons in the same dataframe for input to the CKMR model. Note that not all columns are necessary for each model: for instance, BI is only used when the model accounts for intermittent breeding dynamics, while ref.year and pop.growth.yrs only matter if using a population growth model.

| <u>ref.year</u> | <u>all</u> | <u>yes</u> | <u>mort.yrs</u> | <u>type</u> | <u>parent</u> | <u>ref.year.gap</u> | <u>BI</u> |
| --- | --- | --- | --- | --- | --- | --- | --- |
| 90 | 90 | 0 | 0 | PO | mother | 10 | NA |
| 90 | 449 | 2 | 1 | HS | mother | 10 | off |
| 90 | 90 | 0 | 1 | PO | mother | 10 | NA |
| 90 | 717 | 0 | 2 | HS | mother | 10 | on |
| 90 | 72 | 0 | 2 | PO | mother | 10 | NA |
| 90 | 899 | 1 | 3 | HS | mother | 10 | off |
| 90 | 108 | 1 | 3 | PO | mother | 10 | NA |
| 90 | 845 | 0 | 4 | HS | mother | 10 | on |
| 90 | 611 | 0 | 5 | HS | mother | 10 | off |
| 90 | 537 | 1 | 6 | HS | mother | 10 | on |
| 90 | 378 | 0 | 7 | HS | mother | 10 | off |
| 90 | 234 | 0 | 8 | HS | mother | 10 | on |
| 89 | 100 | 0 | 0 | PO | mother | 9 | NA |
| 89 | 125 | 2 | 0 | PO | mother | 9 | NA |

**Table S3:**
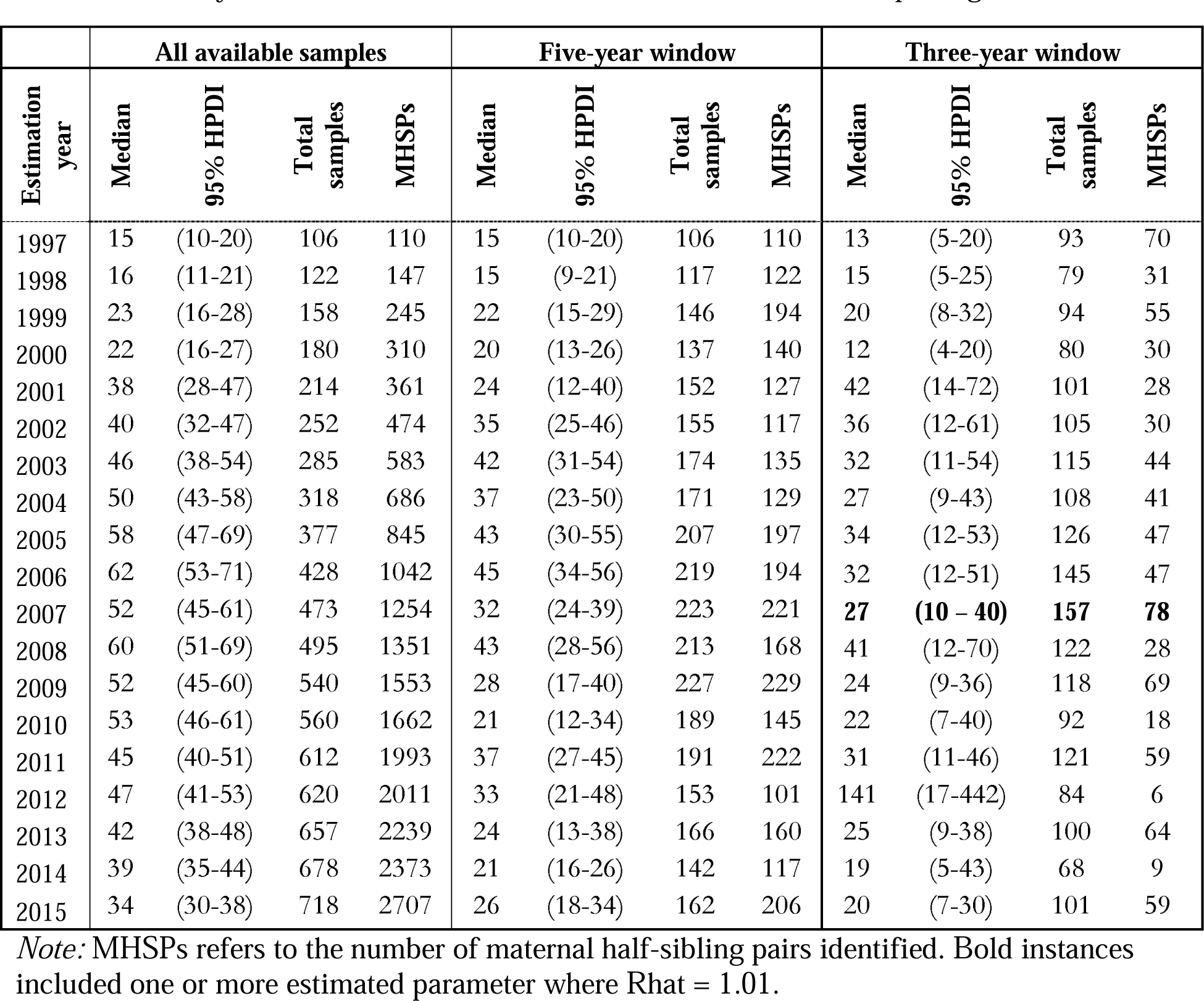
Year-specific abundance estimates for *N*_♀*b(t)*_ from the real dataset for Bimini lemon sharks when only one individual was retained from each mother/father pairing.

**Table S4:**
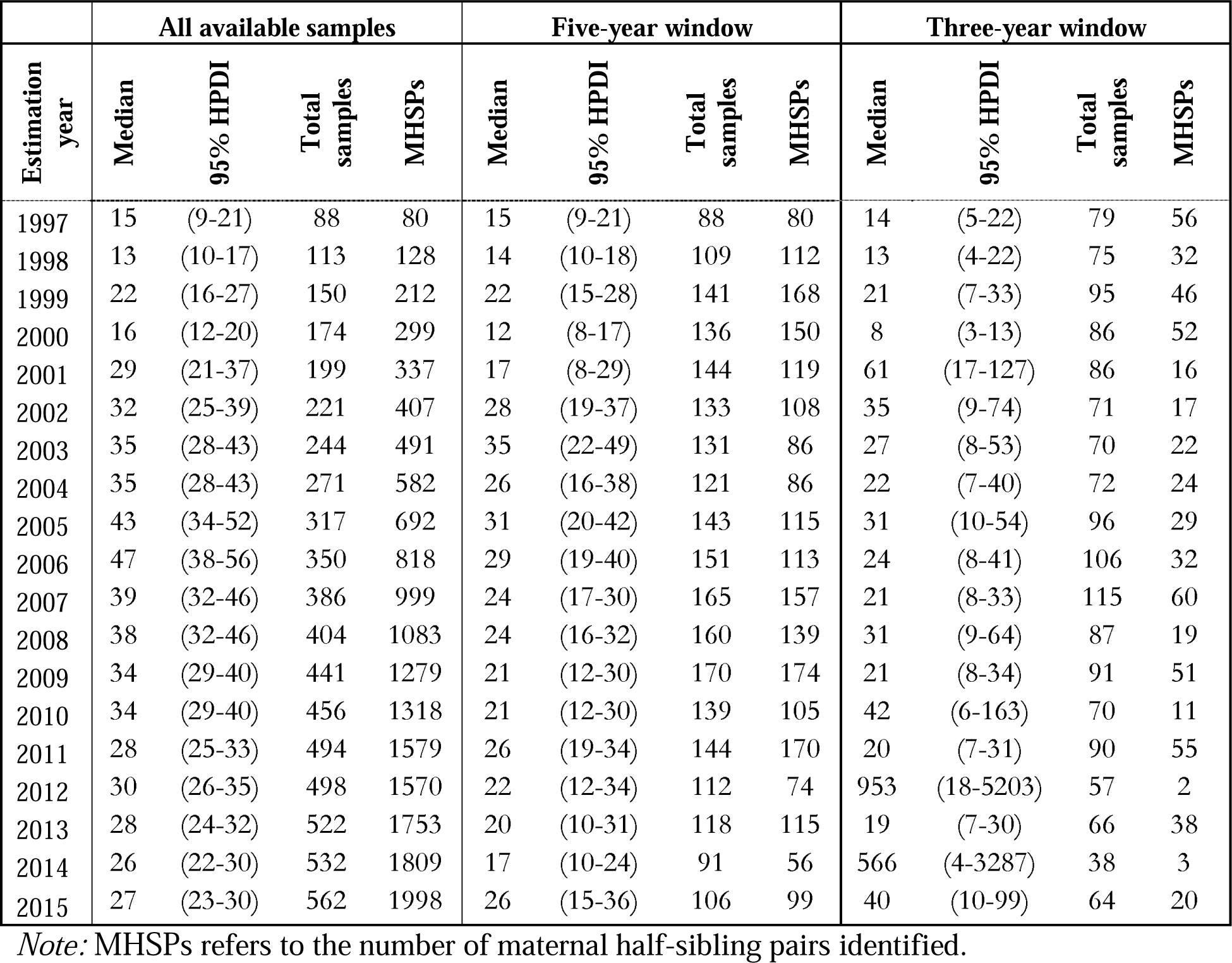
Year-specific abundance estimates for *N*_♀*b(t)*_ from the real dataset for Bimini lemon sharks when the dataset was downsampled to 30% of the full dataset and all individuals were kept from each mother/father pairing. The numbers reported here represent the means over 50 iterations of random downsampling.

| Estimation year | All available samples |  |  |  | Five-year window |  |  |  | Three-year window |  |  |  |
| --- | --- | --- | --- | --- | --- | --- | --- | --- | --- | --- | --- | --- |
|  | Median | 95% HPDI | Total samples | MHSPs | Median | 95% HPDI | Total samples | MHSPs | Median | 95% HPDI | Total samples | MHSPs |
| 1997 | 15 | (9-21) | 88 | 80 | 15 | (9-21) | 88 | 80 | 14 | (5-22) | 79 | 56 |
| 1998 | 13 | (10-17) | 113 | 128 | 14 | (10-18) | 109 | 112 | 13 | (4-22) | 75 | 32 |
| 1999 | 22 | (16-27) | 150 | 212 | 22 | (15-28) | 141 | 168 | 21 | (7-33) | 95 | 46 |
| 2000 | 16 | (12-20) | 174 | 299 | 12 | (8-17) | 136 | 150 | 8 | (3-13) | 86 | 52 |
| 2001 | 29 | (21-37) | 199 | 337 | 17 | (8-29) | 144 | 119 | 61 | (17-127) | 86 | 16 |
| 2002 | 32 | (25-39) | 221 | 407 | 28 | (19-37) | 133 | 108 | 35 | (9-74) | 71 | 17 |
| 2003 | 35 | (28-43) | 244 | 491 | 35 | (22-49) | 131 | 86 | 27 | (8-53) | 70 | 22 |
| 2004 | 35 | (28-43) | 271 | 582 | 26 | (16-38) | 121 | 86 | 22 | (7-40) | 72 | 24 |
| 2005 | 43 | (34-52) | 317 | 692 | 31 | (20-42) | 143 | 115 | 31 | (10-54) | 96 | 29 |
| 2006 | 47 | (38-56) | 350 | 818 | 29 | (19-40) | 151 | 113 | 24 | (8-41) | 106 | 32 |
| 2007 | 39 | (32-46) | 386 | 999 | 24 | (17-30) | 165 | 157 | 21 | (8-33) | 115 | 60 |
| 2008 | 38 | (32-46) | 404 | 1083 | 24 | (16-32) | 160 | 139 | 31 | (9-64) | 87 | 19 |
| 2009 | 34 | (29-40) | 441 | 1279 | 21 | (12-30) | 170 | 174 | 21 | (8-34) | 91 | 51 |
| 2010 | 34 | (29-40) | 456 | 1318 | 21 | (12-30) | 139 | 105 | 42 | (6-163) | 70 | 11 |
| 2011 | 28 | (25-33) | 494 | 1579 | 26 | (19-34) | 144 | 170 | 20 | (7-31) | 90 | 55 |
| 2012 | 30 | (26-35) | 498 | 1570 | 22 | (12-34) | 112 | 74 | 953 | (18-5203) | 57 | 2 |
| 2013 | 28 | (24-32) | 522 | 1753 | 20 | (10-31) | 118 | 115 | 19 | (7-30) | 66 | 38 |
| 2014 | 26 | (22-30) | 532 | 1809 | 17 | (10-24) | 91 | 56 | 566 | (4-3287) | 38 | 3 |
| 2015 | 27 | (23-30) | 562 | 1998 | 26 | (15-36) | 106 | 99 | 40 | (10-99) | 64 | 20 |

*Note:* MHSPs refers to the number of maternal half-sibling pairs identified.

**Table S5:** Year-specific abundance estimates for *N*_♀*b(t)*_ from the real dataset for Bimini lemon sharks when the dataset was downsampled to 30% of the full dataset and only one individual was retained from each mother/father pairing. The numbers reported here represent the means over 50 iterations of random downsampling.

| Estimation year | All available samples |  |  |  | Five-year window |  |  |  | Three-year window |  |  |  |
| --- | --- | --- | --- | --- | --- | --- | --- | --- | --- | --- | --- | --- |
|  | Median | 95% HPDI | Total samples | MHSPs | Median | 95% HPDI | Total samples | MHSPs | Median | 95% HPDI | Total samples | MHSPs |
| 1997 | 14 | (9-22) | 60 | 37 | 14 | (9-22) | 60 | 37 | 18 | (10-29) | 54 | 25 |
| 1998 | 13 | (8-21) | 72 | 53 | 16 | (8-28) | 70 | 45 | 39 | (11-102) | 47 | 11 |
| 1999 | 21 | (14-30) | 94 | 91 | 22 | (13-34) | 88 | 73 | 30 | (15-54) | 57 | 21 |
| 2000 | 17 | (11-24) | 106 | 116 | 17 | (9-27) | 82 | 55 | 16 | (7-32) | 47 | 12 |
| 2001 | 37 | (25-51) | 123 | 129 | 40 | (21-63) | 87 | 48 | 93 | (26-260) | 53 | 8 |
| 2002 | 34 | (24-46) | 141 | 162 | 30 | (16-49) | 83 | 40 | 93 | (19-387) | 49 | 8 |
| 2003 | 36 | (27-46) | 155 | 200 | 38 | (20-65) | 87 | 39 | 52 | (17-129) | 53 | 11 |
| 2004 | 39 | (29-49) | 171 | 223 | 39 | (19-70) | 82 | 35 | 59 | (18-139) | 51 | 11 |
| 2005 | 52 | (41-64) | 202 | 281 | 51 | (25-86) | 100 | 56 | 72 | (27-143) | 65 | 15 |
| 2006 | 58 | (47-71) | 229 | 333 | 70 | (32-119) | 108 | 52 | 61 | (23-127) | 74 | 15 |
| 2007 | 58 | (47-69) | 253 | 398 | 49 | (25-81) | 114 | 71 | 41 | (19-77) | 82 | 26 |
| 2008 | 56 | (46-67) | 265 | 433 | 41 | (20-67) | 109 | 55 | 54 | (17-135) | 61 | 9 |
| 2009 | 50 | (42-59) | 288 | 495 | 22 | (16-30) | 118 | 74 | 29 | (17-47) | 63 | 21 |
| 2010 | 52 | (43-61) | 299 | 535 | 21 | (14-29) | 97 | 44 | 41 | (8-201) | 48 | 7 |
| 2011 | 43 | (37-49) | 323 | 618 | 21 | (16-27) | 100 | 74 | 31 | (18-50) | 63 | 21 |
| 2012 | 43 | (37-50) | 332 | 654 | 20 | (12-31) | 79 | 33 | 1304 | (20-6180) | 43 | 2 |
| 2013 | 35 | (30-40) | 350 | 733 | 20 | (14-27) | 85 | 52 | 25 | (14-42) | 50 | 19 |
| 2014 | 35 | (30-40) | 363 | 790 | 15 | (10-23) | 68 | 28 | 825 | (9-4807) | 30 | 2 |
| 2015 | 31 | (27-35) | 372 | 839 | 19 | (13-26) | 79 | 51 | 38 | (16-81) | 48 | 13 |

*Note:* MHSPs refers to the number of maternal half-sibling pairs identified.

**Figure S1:**
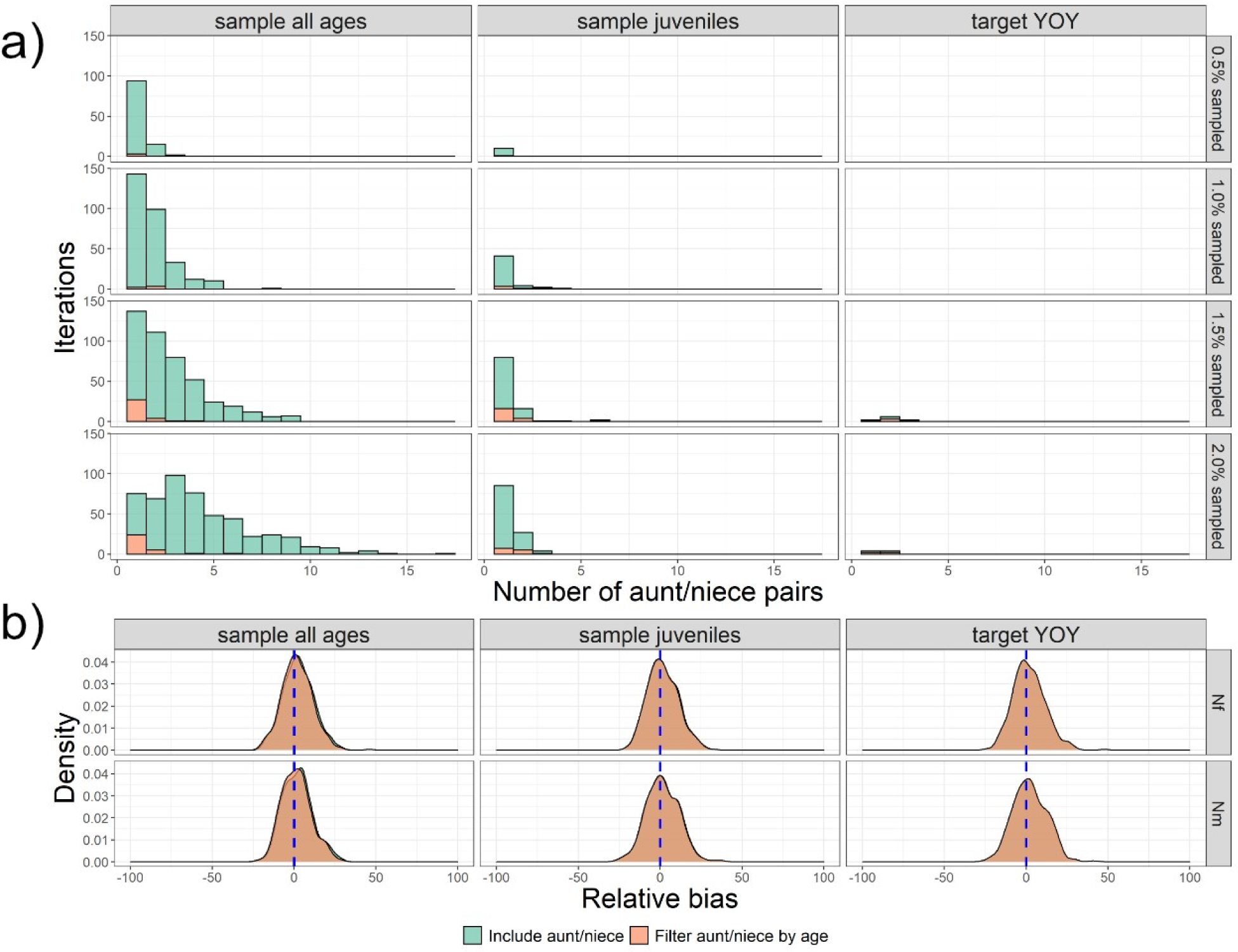
Prevalence of aunt/niece (uncle/nephew, etc.) pairs and effects on model performance by sampling scheme and sampling intensity. **a)** Histogram summarizing the number of identified aunt/niece (uncle/nephew, etc.) pairs. The Y axis represents the number of iterations that contained the number of aunt/niece pairs specified by the X axis. Green represents the total number of aunt/niece etc. pairs identified; orange represents the number of pairs remaining after instituting a filter that removed pairwise comparisons with a year gap greater than the age of reproduction. Most simulations contained 0 aunt/niece pairs, so these were removed for visualization purposes. **b)** Effects of mistakenly including aunt/niece pairs as half-siblings before (green) and after (orange) a year gap filter when 2% of the population was sampled. The effect on bias was minimal, and the distributions overlapped almost perfectly (though there is a very slight observable difference for the sample all ages scenario).

**Figure S2:**
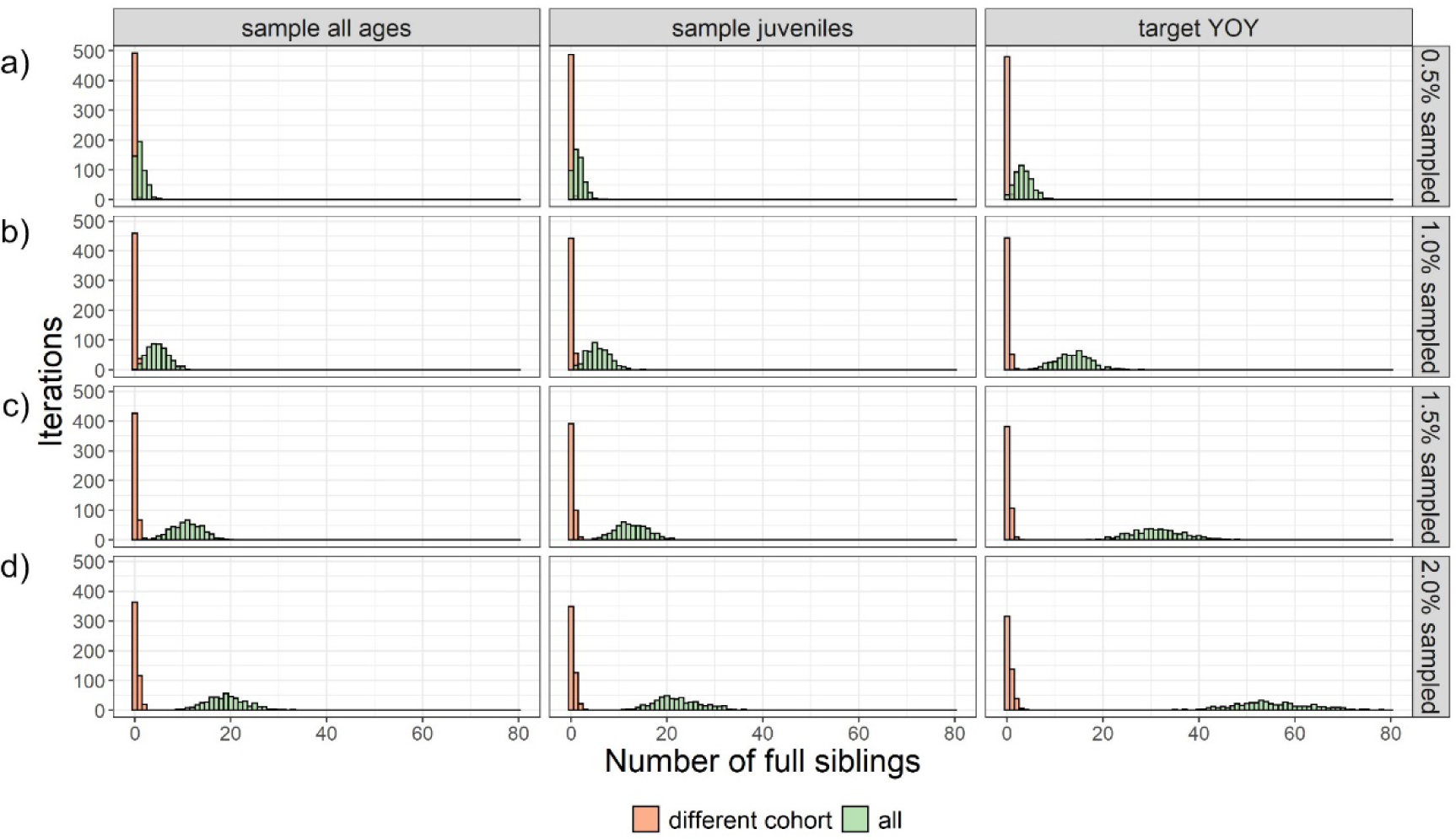
Histogram of full siblings observed for different sampling schemes and intensities, collated over 500 iterations. The Y axis represents the number of iterations that contained the number of full sibling pairs specified by the X axis The total number of full siblings (green) includes within-cohort full siblings and represents almost all instances. The number of cross-cohort full siblings (orange) was substantially lower.

**Figure S3:**
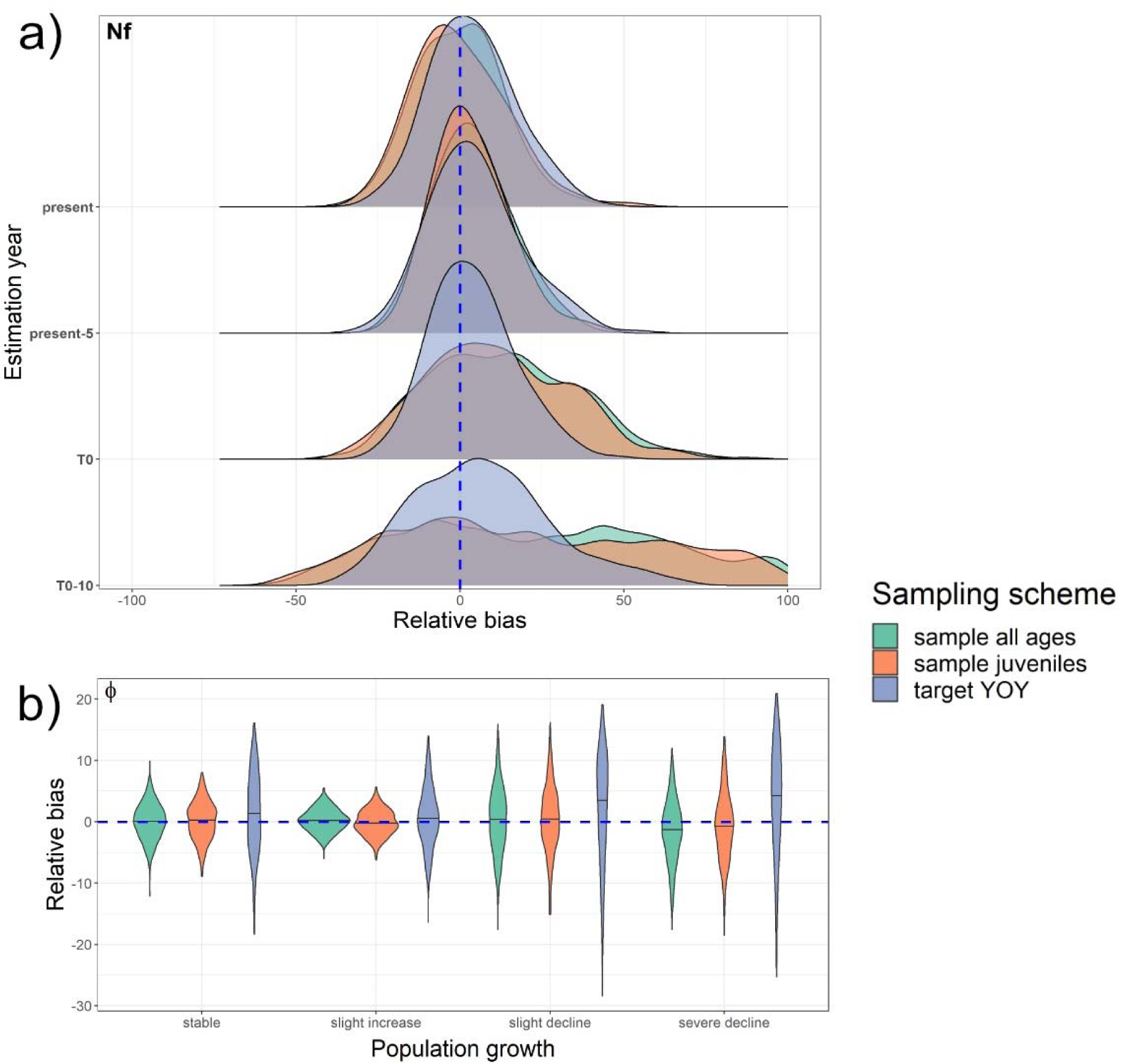
Estimates of **a)** female abundance (*Nf*) when a parameter for population growth (λ) was included in the model and realized population growth was stable, and **b)** survival (L) under different population growth scenarios.

**Figure S4:**
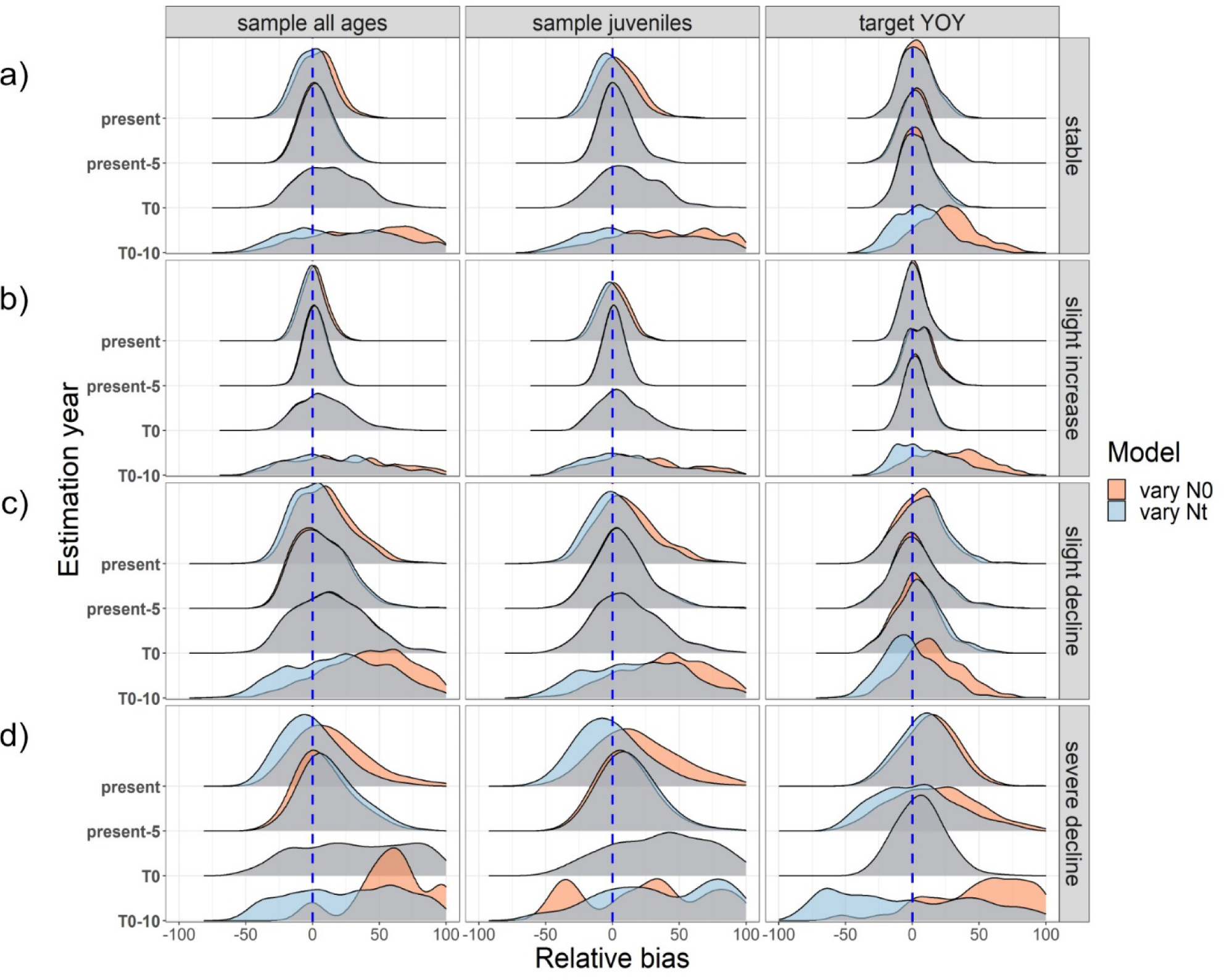
Comparison of relative bias for different population growth scenarios and approaches to integrating a population growth model. The orange density plots come from a model where abundance in year *t* was directly estimated in year *t* (i.e., the “Estimation year”) using Eqs. 4 and 5. The blue density plots come from the primary model we used throughout our simulations, where *t_0_* represented the first instance of *y_j_* in each dataset, and abundance in year *t* was derived from estimates of *N*_♀*(t0)*_ using Eq. 3.

**Figure S5:**
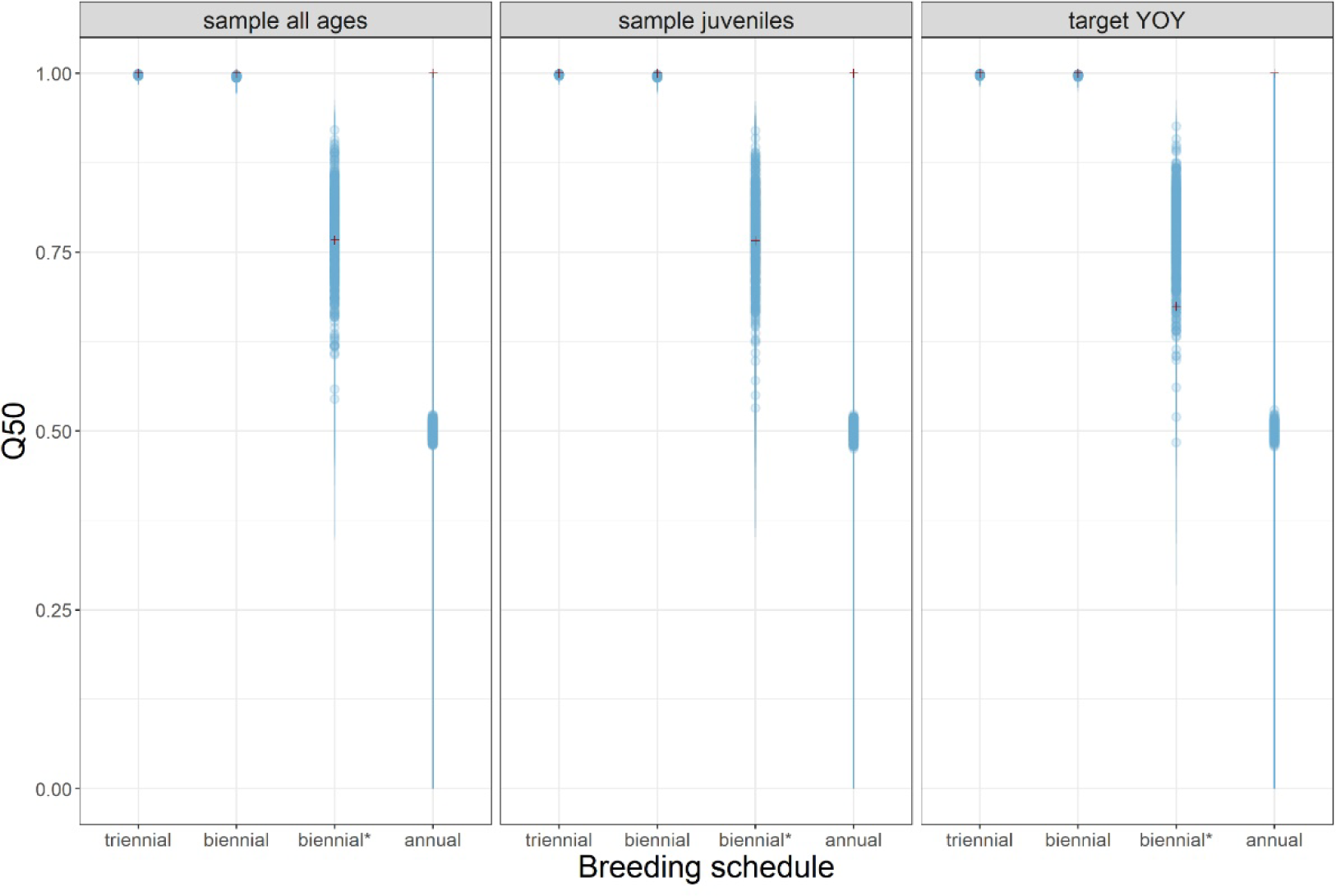
Estimates of ψ for multiennial breeding scenarios. The blue points represent the distribution of estimates, and the red crosses represent the mean realized value of psi across all 500 iterations, calculated as the proportion of total positive comparisons that came from on-cycle breeders.

**Figure S6:**
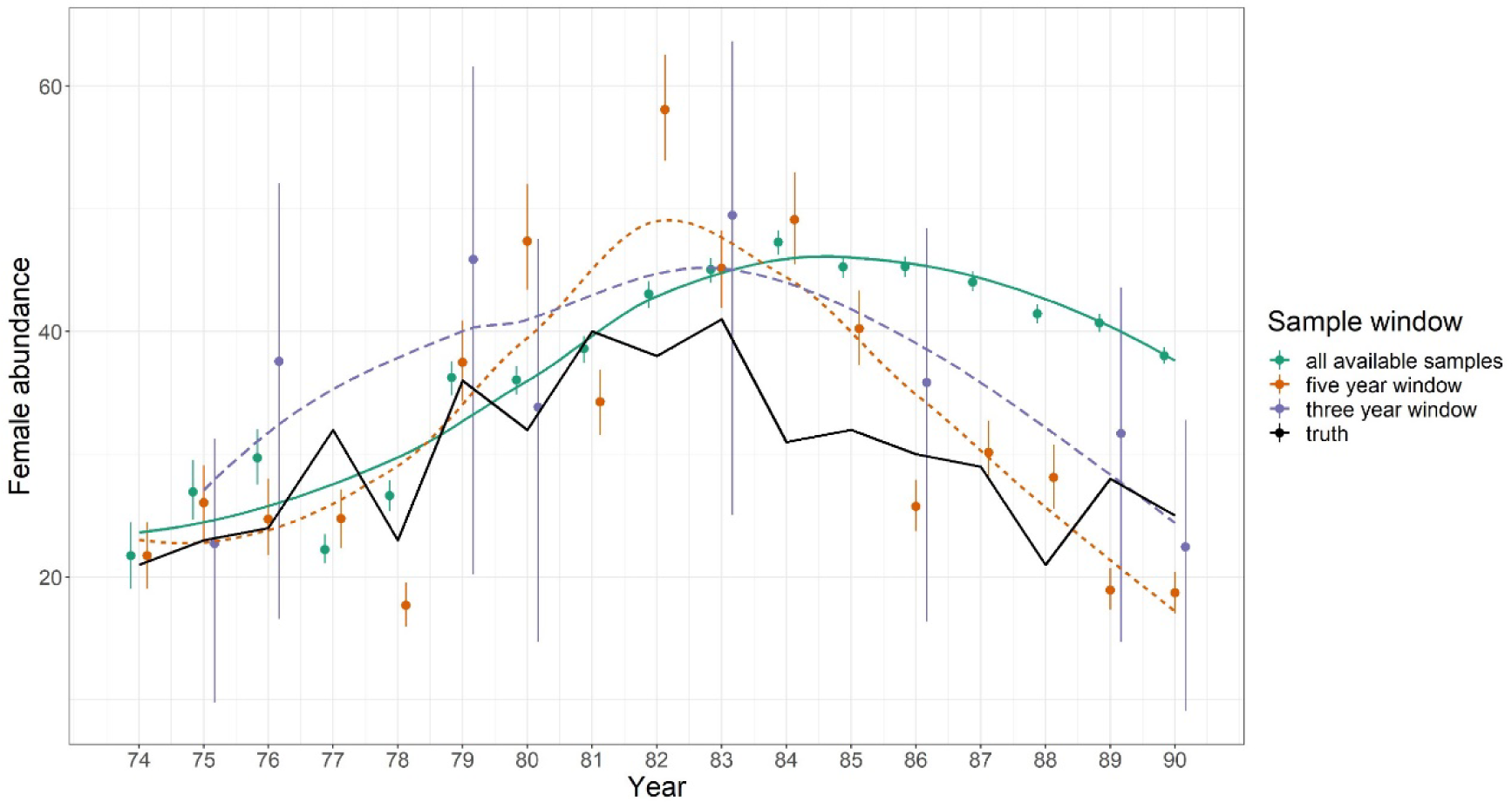
A representative iteration of a time-series of abundance estimates for females from Bimini lemon shark simulations when full siblings are retained in the analysis. Estimates for each year for three different sampling windows are shown relative to the truth (black). Trends are visualized using a loess regression.

**Figure S7:**
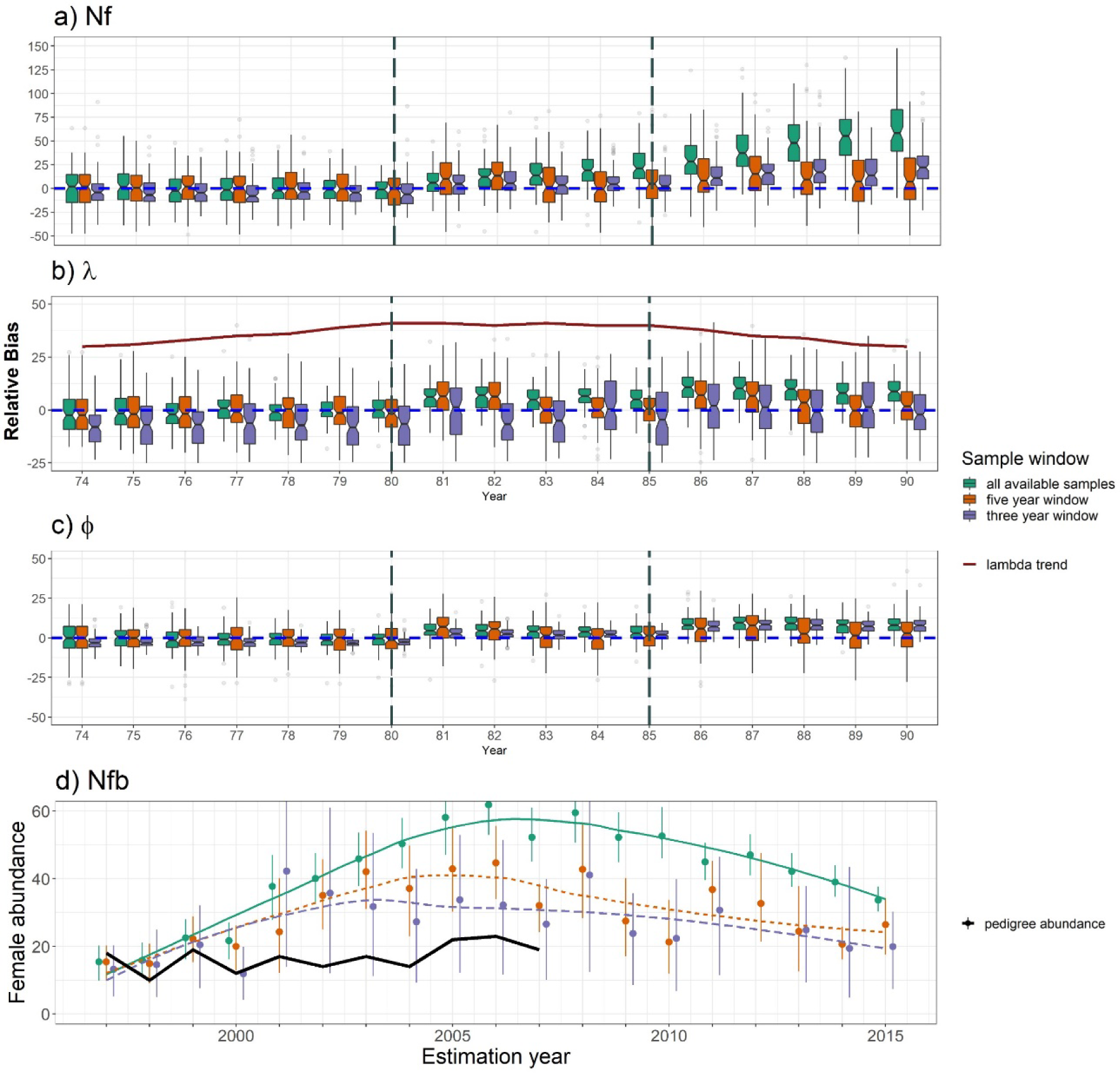
Time series of CKMR parameter estimates for simulated (**a-c**) and real (**d**) female lemon sharks at Bimini, Bahamas when only one individual from each mother/father pairing is retained from each dataset. The colors and lines are the same as Figure 6. **a-c)** Relative bias from 100 distinct population simulations and model fits. **a)** Relative bias of abundance estimates for adult females (*Nf*) in each year of the time series. **b)** Relative bias of λ estimates relative to the observed population growth rate in the associated estimation year. **c)** Relative bias of survival (□) relative to the observed survival rate in the associated estimation year. **d)** Abundance estimates for breeding females (*Nfb*), derived from estimates of total *Nf* using Eq.6, in the North Bimini Lagoon using real genetic data. Points represent the median of the posterior distribution, and error bars reflect the 95% highest posterior density interval (HPDI). The trend is visualized using a loess regression. The black line labeled as “pedigree abundance” is a time-series of abundance estimates for the population that was independently derived for the population by Dibattista et. al. (2011).

**Figure S8:**
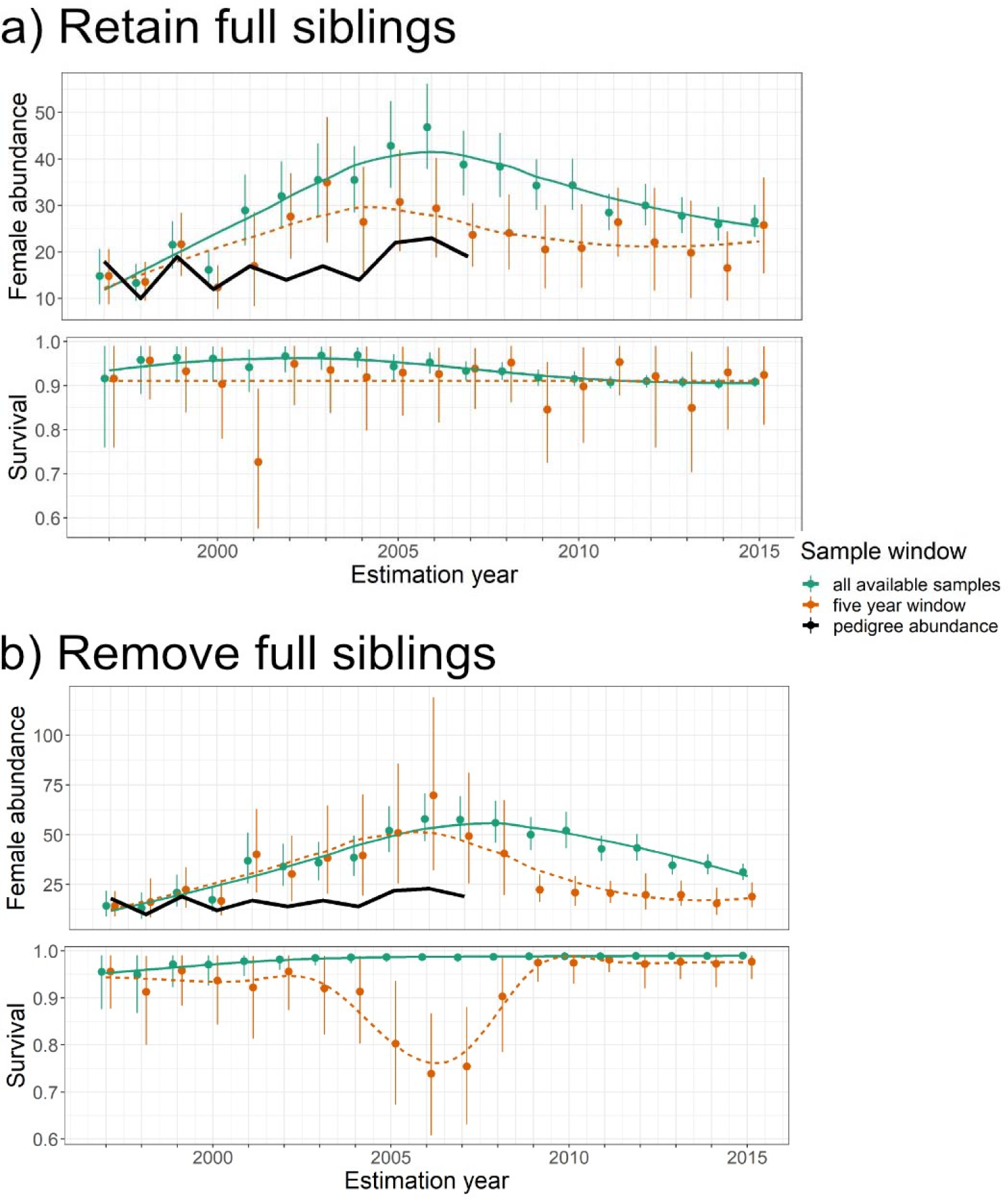
Estimates of abundance and survival for females in the North Bimini Lagoon using datasets that were downsampled and three different sampling windows. Estimates represent the means over 50 iterations of random downsampling. **a)** Estimates of abundance and survival for breeding females using 30% of the total samples for each year without removing full siblings. **b)** Estimates of abundance and survival for breeding females using 30% of the total samples for each year and also removing full siblings.

**Figure S9:**
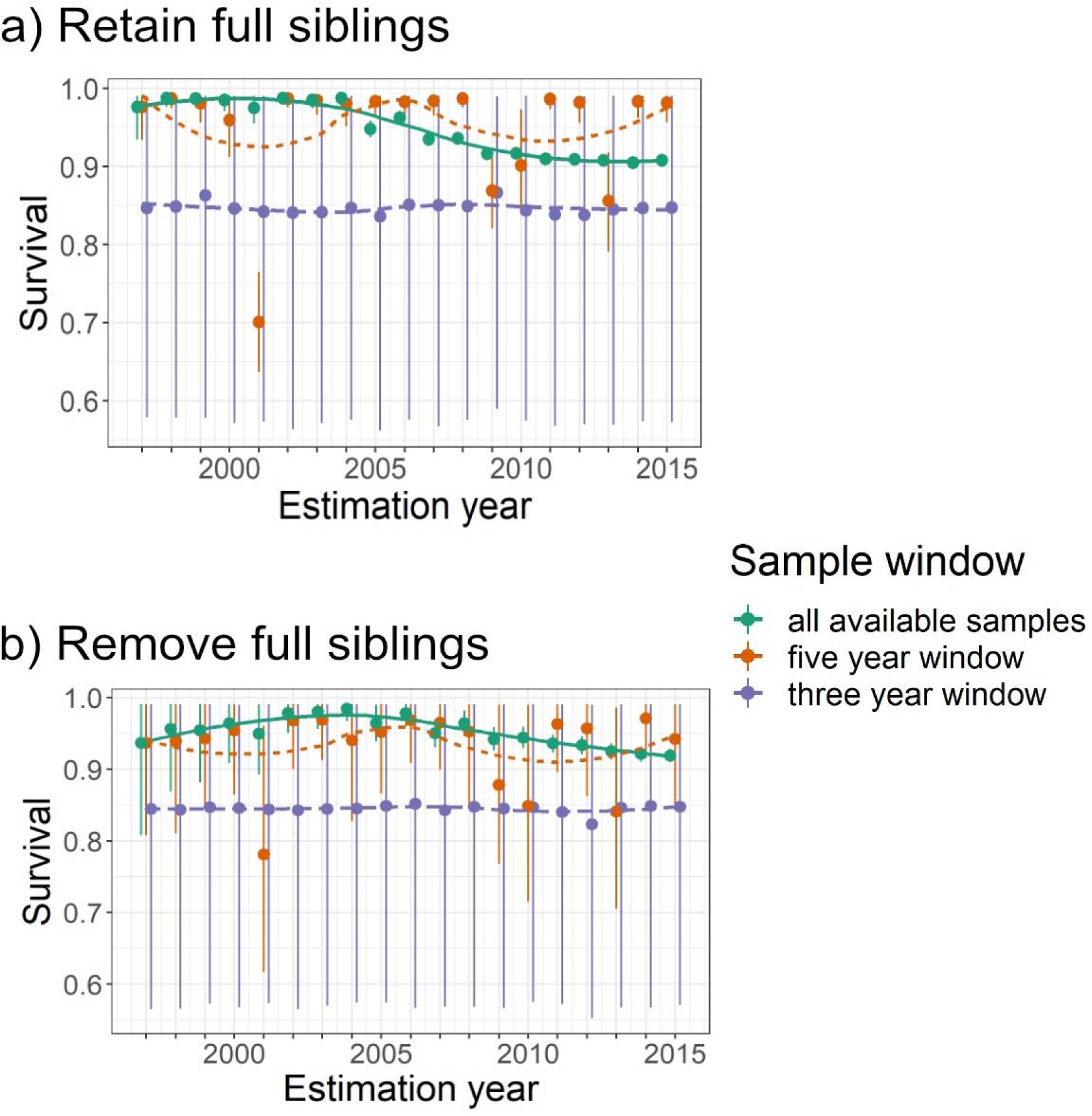
CKMR-based survival estimates for Bimini lemon sharks using real data and three different sampling windows when **a)** full siblings were retained in the dataset and **b)** when full siblings were removed from the dataset. Trends were visualized using a loess regression.

**Figure S10:**
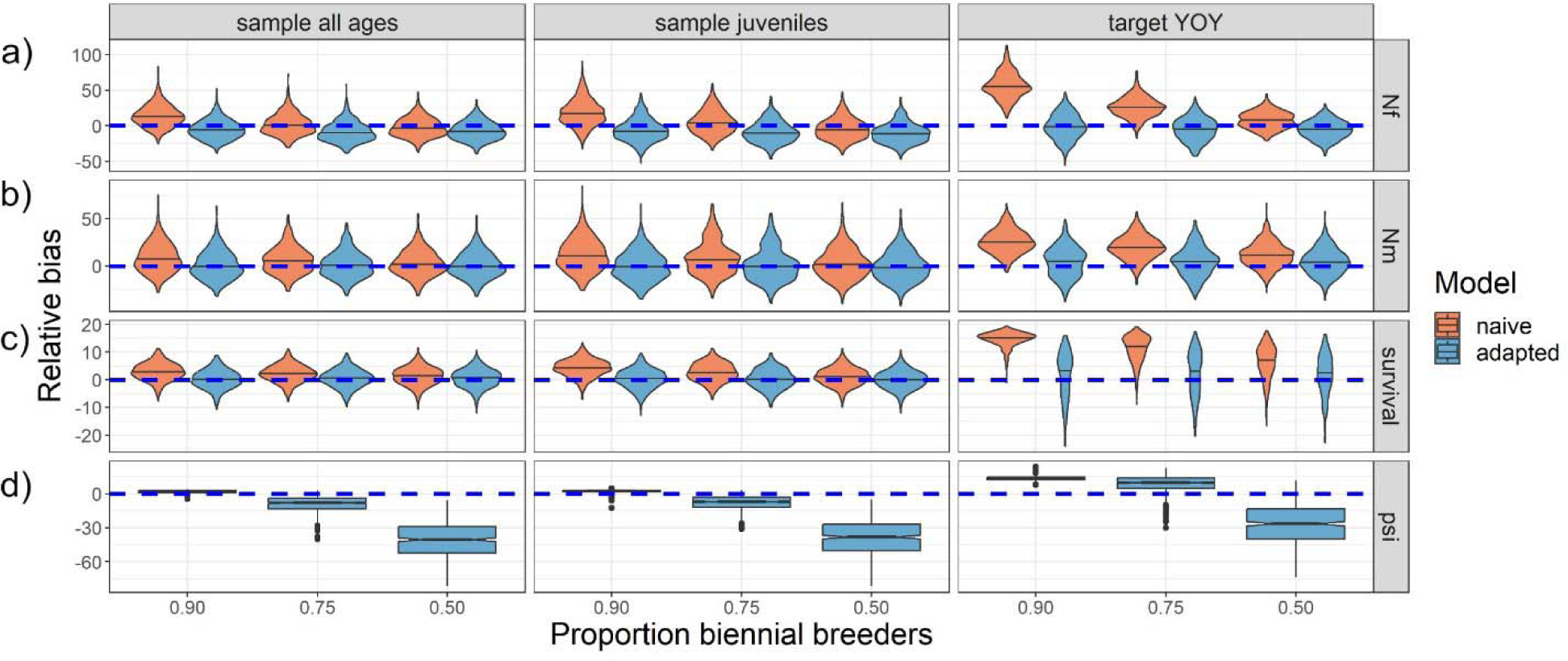
Relative bias of parameter estimates in a simulated population with different ratios of biennial vs. annual female breeders, holding fecundity constant such that lifetime reproductive output varied among the different portions of the population. Both annual and biennial models were fit to the data. **a)** Relative bias of abundance estimates for females. **b)** Relative bias of abundance estimates for males. Note that males bred annually, but shared parameters for survival (□) and population growth (λ) with females. **c)** Relative bias of shared survival estimates. **d)** Relative bias of estimates of ψ with different proportions of biennial breeders.

